# Creating a ‘Timeline’ of ductal carcinoma in situ to identify processes and biomarkers for progression towards invasive ductal carcinoma

**DOI:** 10.1101/2022.03.01.482529

**Authors:** Clare A. Rebbeck, Jian Xian, Susanne Bornelöv, Joseph Geradts, Amy Hobeika, Heather Geiger, Jose Franco Alvarez, Elena Rozhkova, Ashley Nicholls, Nicolas Robine, Herbert K. Lyerly, Gregory J. Hannon

## Abstract

Ductal carcinoma in situ (DCIS) is considered a non-invasive precursor to breast cancer, and although associated with an increased risk of developing invasive disease, many women with DCIS will never progress beyond their in situ diagnosis. The path from normal duct to invasive disease is not well understood, and efforts to do so are hampered by the substantial heterogeneity that exists between patients and even within patients. Using gene expression analysis, we have generated a ‘Timeline’ of disease progression, utilising the variability within patients and combining >2,000 individually micro-dissected ductal lesions from 145 patients into one continuous trajectory. Using this Timeline we show there is a progressive loss in basal layer integrity, coupled with two epithelial to mesenchymal transitions (EMT), one early in the timeline and a second just prior to cells leaving the duct. We identify early processes and potential biomarkers, including *CAMK2N1*, *MNX1*, *ADCY5*, *HOXC11* and *ANKRD22*, whose reduced expression is associated with the progression of DCIS to invasive breast cancer.

## Introduction

Ductal carcinoma in situ (DCIS) is considered to be a non-invasive precursor to breast cancer, and when found is associated with an approximate 10-fold increased risk of developing an invasive carcinoma ^1^. However, over half of untreated DCIS patients may never develop breast cancer ^2,3^. Despite this, conventional treatment typically comprises either mastectomy or breast conserving surgery coupled with radiation. In order to treat women most effectively and reduce unnecessary treatment, it is vital that we understand more about DCIS and what factors influence the risk of progression to invasive disease. At present, the path from normal ductal epithelium to invasive ductal carcinoma (IDC) remains poorly understood. Current thinking suggests that there is a step-wise progression from a normal duct, through atypical ductal hyperplasia (ADH), to DCIS followed by microinvasion from the duct to established invasive ductal carcinoma. While the ductal epithelium is typically comprised of a mixture of luminal and basal-like cells, ADH and DCIS are expansions of the luminal compartment ^4^, with the presence of nuclear and/or architectural atypia. Distinguishing DCIS from ADH is one of the most difficult challenges in breast pathology, and there is marked inter-observer variability, suggesting that not all disease states are easily categorized by morphology alone. In addition, distinguishing low-risk from high-risk DCIS lesions can be difficult at best. A number of studies have examined transcriptional differences between normal ductal tissue, ADH, DCIS, and IDC ^5-8^; however, there has been little agreement surrounding genes that mark transitions between tissue states, and studies have often been limited by patient number and tissue quality.

Here we describe the analyses of a large-scale transcriptomic study of over 2700 pathologically annotated and individually micro-dissected regions from 145 fresh-frozen patient biopsies. Focusing largely on DCIS, we combined 1624 RNAseq libraries from DCIS with 394 libraries from IDC, 258 from atypical ductal lesions, 237 from benign ductal lesions and a further 211 libraries from normal mammary epithelium. Using this data, we were able to describe the evolution of tissue states from the transcriptional changes characteristic of very early lesions, through progression toward, and development of invasive carcinoma. This pseudo-‘timeline’ of disease progression revealed processes characteristic of different points along the path from normal epithelium to IDC. Considering both Pure DCIS (where no IDC was found in that patient, nor diagnosed from that patient during 10+ years of care) and DCIS from patients diagnosed with co-occurring IDC, we saw that the position of individual lesions on the timeline was not dictated solely by patient diagnosis. Even among lesions derived from patients having only DCIS, there existed a range of developmental stages, as defined along our timeline, that mirrored those seen in patients that progressed to IDC. We also found that position along the timeline was not determined by ER/PR or Her2 status similar to a prior finding detailing a trajectory of changes surrounding tumour stroma^9^, thus potentially indicating that early stage disease results from changes in the same core processes for both of ER+ and ER-negative lesions.

## Results

### The cohort

145 Frozen tissue biopsies, kindly donated to Duke University, were microdissected for DCIS, IDC and other ductal regions of interest (see methods for further details). Each individual lesion was carried through for RNA sequencing and quality control checks (see Supplementary methods). We found that 68% of patients had DCIS mRNA expression patterns that matched their clinical scoring for estrogen receptor (ER/ *ESR1*), progesterone receptor (PR/ *PGR*), and human epidermal growth factor receptor 2 (Her2/ *ERBB2*), (Table S4). Of 44 patients (32%) that did not, 6 showed a clear difference in ER status, 29 showed a clear difference in PR status, and 8 showed a clear difference in Her2 status (where Her2 had been clinically scored). It must be acknowledged, however, that where IDC was found in the clinical diagnostic biopsy, it is the IDC that was scored for these markers and not the DCIS, and scoring is based on a number of factors, such as the percentage of invasive tumour cells with nuclear staining as well as the average staining intensity. Within the IDC samples, we also found that 68% of patients matched their clinical scoring for ER, PR, and Her2, and the remaining 32% (13 patients) showed distinct deviations in their RNA expression from that of the clinical scoring. Four patients had a clear difference in RNA expression signatures for *ESR1*, *PGR* and *ERBB2*, between their DCIS and IDC. These findings are consistent with the well-established heterogeneity within this disease, and in some cases, we found that different DCIS samples scored differently even within the same tissue section, most often for PGR.

### Triple-Negative DCIS cluster separately

To assess whether there were any distinct groups of DCIS samples, we carried out Principal component analysis (PCA) followed by uniform manifold approximation and projection (UMAP) using only DCIS samples (Fig. 1A) This revealed that the majority of samples largely group together, however Basal-like (as defined by AIMS^10^) triple negative (TN) DCIS samples, with low expression for *ESR1* (ER), *PGR* (PR) and *ERBB2* (Her2) (Fig. 1B), form a distinct cluster away from other DCIS samples, including other non-basal-like TN DCIS samples (Fig. 1A and see fig. S1 for all sample subtype classifications by patients). This is in line with a recent study looking at DCIS subtypes ^11^. Differential expression analysis between this Basal-like TN cluster against the other clusters revealed that the pioneer factor, Forkhead Box A1, *FOXA1* and Melanophilin*, MLPH* mRNA levels were significantly reduced in this cluster as compared to the other subtypes. Other genes showing a strong association with this group are Carbonic anhydrase 12 (*CA12*), transcription factors *SPDEF, FOXC1,* and *ELF5*, sodium channel epithelial subunit, *SCNN1A*, and pyridoxyl kinase, *PDXK* (Fig. 1C). *FOXA1* and*, MLPH* are among other genes annotated as being more highly expressed in luminal cells compared to basal cells, and vice-versa for *ELF5*, in studies of mouse mammary glands ^12^.

**Fig. 1.**
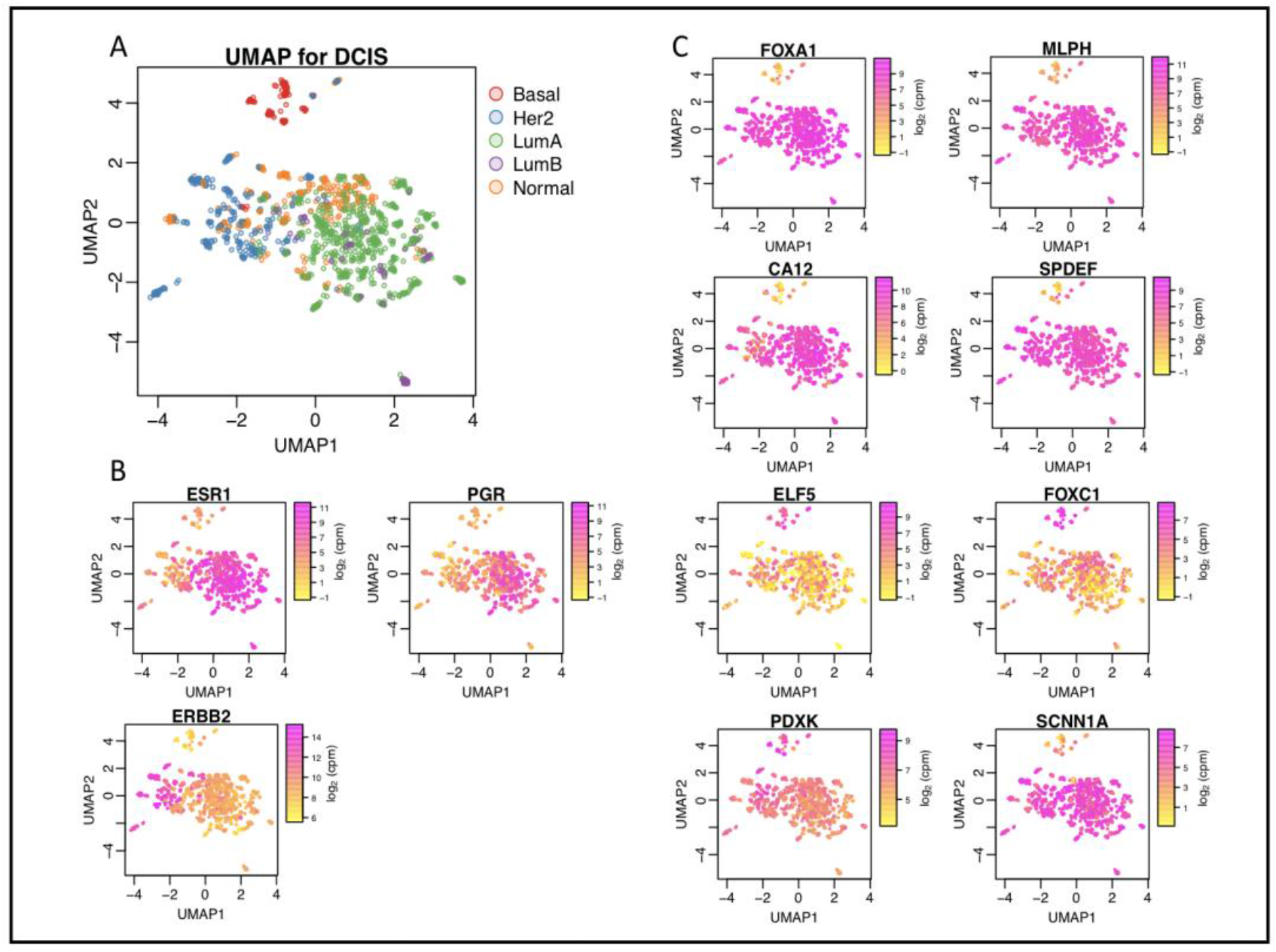
Triple negative DCIS has a transcriptome distinct from other DCIS subtypes. UMAP plots illustrating expression patterns in 1414 DCIS samples by (**A**) AIMS subtype, (**B**) *ESR1*/*PGR*/*ERBB2* gene expression, and (**C**) expression of genes that correlate with triple negative status in DCIS.

*FOXA1* has recently been highlighted as a potentially useful marker for triple negative breast cancer ^13^, and its expression has been suggested to act as a repressor for a subset of basal signature genes ^14^. The association of FOXA1 and triple-negative status has not previously been examined in DCIS, and reports thus far have dismissed a role for FOXA1 as a subtype marker for DCIS as no correlation could be seen with protein expression and that of ER ^15,16^. Here we also observed that *FOXA1* expression does not systematically differ between ER+ and ER-samples, and its reduced expression is only associated with the basal-like TN samples. The substantial overlap between TN-associated markers identified here, and those found by other studies on invasive breast cancer (including *MLPH*, *CA12*, *FOXA1*, *SPDEF*, *FOXC1*), suggest there is a clear distinction of this subtype even at the pre-invasive stage ^17-19^.

### Two gene networks dominate expression differences between co-occurring DCIS and early invasive breast cancer

We sought to leverage our extensive datasets to identify transcriptional differences between DCIS and co-occurring IDC. This would act as a starting point in identifying genes that may be related to the progression towards IDC. We compared the two tissue types from DCIS + IDC patients, (only for those where we had useable data for both tissue types within a patient), N = 33. Using this criterion, we aimed to compare samples that were most closely matched to minimise inherent inter-patient variability. We carried out differential expression analysis between the two tissue groups and found 401 significantly differentially expressed genes (DEGs). Taking the 53 genes with an Adj. P value < 0.00001, we used STRING ^20^ to examine their connectivity. We were surprised to find that the genes formed two main, highly interconnected networks, with very few unconnected genes (Fig. 2A). Gene Ontology (GO) term analysis on these two networks revealed an enrichment for upregulated (in IDC over DCIS) genes involved in both extracellular matrix (EM) organisation (FDR 2.5E-16, Fold Enrichment; 29) and cell adhesion (FDR 1.8E-6, Fold Enrichment; 7) and down-regulated genes associated with both epidermis development (FDR 5.3E-8, Fold Enrichment; 18) and epithelial development (FDR 1.6E-06, Fold Enrichment; 7). Specific genes included in each cluster network are frequently associated with these processes, such as *FN1* (Fibronectin) and the collagen genes (*COL1A2*, *COL1A1*, *COL12A*, *COL3A1*, *COL5A2*). Other genes, such as *MMP11* (matrix metalloproteinase 11) and *POSTN* (Periostin) are involved in epithelial cell adhesion and migration, and *THBS2* (Thrombospondin 2), a mediator of cell-cell and cell-matrix interactions. The second cluster network includes *DSC3* and *DSG3*, reported to be expressed only in myoepithelial cells within the basal cell layer, *KRT5*, *KRT14*, *KRT6B* and *KRT15*, markers for basal epithelial cells, and *KLK5* and *KLK7*, considered to be involved in desquamation ^21^. We noted a substantial overlap between genes in this cluster and those found to be differentially expressed in basal cells (as compared to luminal cells) in mouse and human mammary glands ^22,23^. Considered together, these data could suggest that the expression changes we observe in the down-regulated genes may be reflective of a loss in the basal compartment of the duct. Carrying out the same differential analysis on a per subtype basis had limited statistical power for all but the Luminal A subtype, due to the reduced sample sizes (Her2 N = 6, LumB N = 3, Basal N = 3, Normal Like N = 3). However, we did find many of the 53 genes from the combined analysis also ranked highly in the individual subtype analyses, with Basal patients sharing the least (DEGs can found in Table S5).

**Fig. 2.**
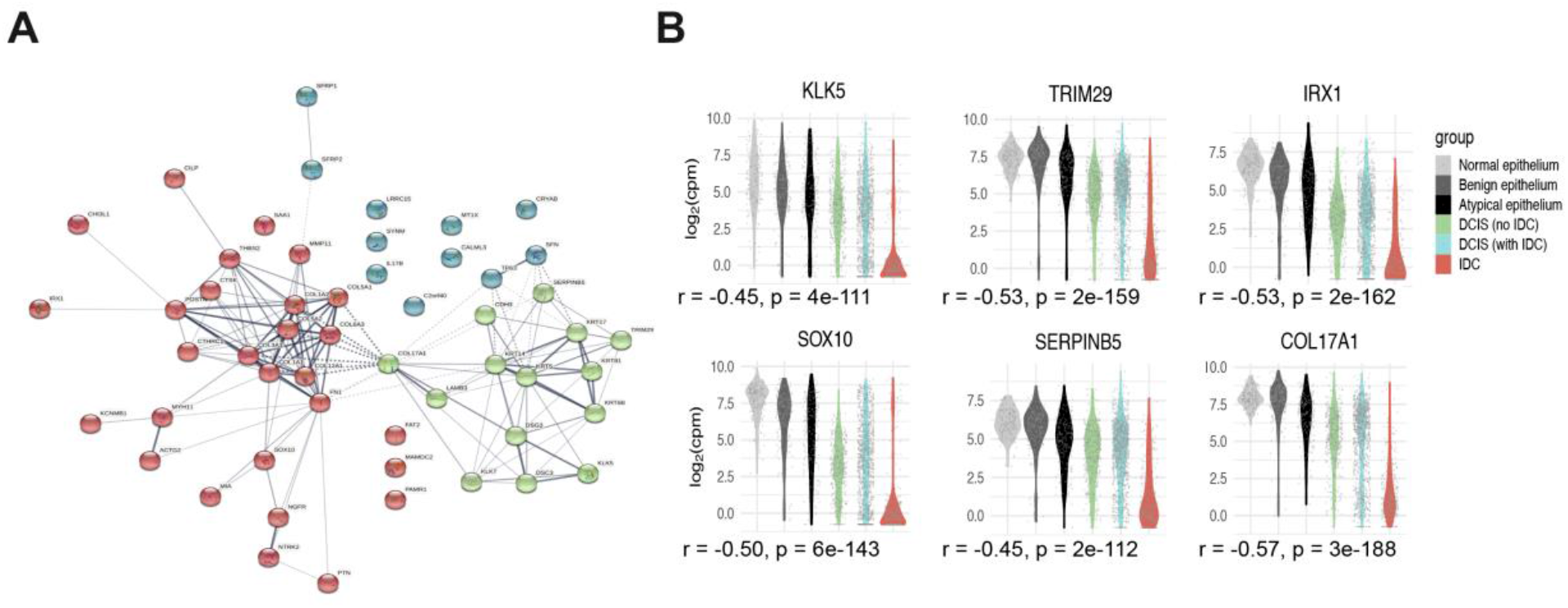
Differentially expressed genes between DCIS and co-occurring IDC. (**A**) String connectivity with k-means clustering [3 clusters] of the top 53 significant genes. (**B**) Expression distribution for example genes that showed a progressive shift among different tissue groups. The Spearman rank correlation (between expression and ordered tissue groups) is given as r = rho.

### A pseudo ‘timeline’ of DCIS progression from normal epithelium to IDC

Given the strong presence of just two dominant processes that appeared to be contributing to the transition from DCIS to IDC, we examined how the integrity of the basal layer and the EM may differ in our other tissue types, or disease statuses (Pure DCIS or Not Pure DCIS). Looking at these same 53 genes we noted in some cases a progressive shift from expression levels in normal ductal tissue to that seen in IDC (Fig. 2B and fig. S3). Interestingly, for some genes, some DCIS samples displayed an expression pattern that was more reflective of normal epithelium while others more closely resembled IDC, even if the samples were isolated from the same patient. This led us to hypothesize that some DCIS samples are more closely related to their normal counterparts and others more related to their invasive counterparts, and that perhaps this indicated a possible continuum of tissue states represented during disease progression, within an individual patient.

**Fig. 3.**
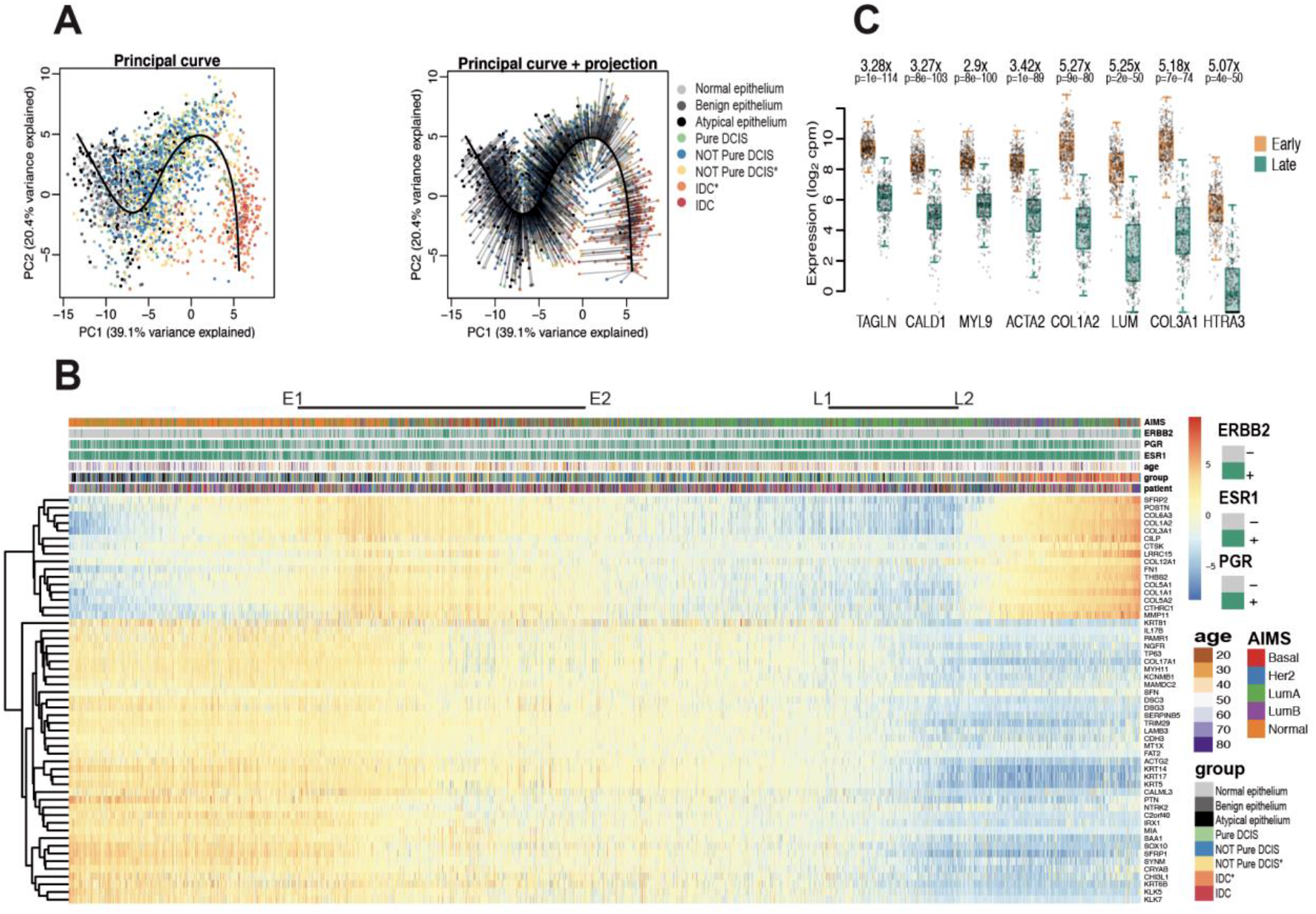
Generating a pseudo-timeline for DCIS. (**A**) PCA plot based on the most significant (p<0.00001) DEGs between DCIS and co-occurring IDC. All samples and their fitted principal curve shown (left), or with their projection onto the curve (right). (**B**) Heatmap showing expression of each of the 53 genes with samples ordered by their projection to the Principal Curve. Top bars indicate AIMS subtype classification, *ERBB2*, *PGR* and *ESR1* status, age of patient at the time of consent, tissue classification group for each sample, and patient distribution. Relative expression is provided as log2 CPM minus the mean log2 CPM for each gene. E1 – E2 indicate the Early stage and L1 – L2 indicates the Late stage. The ‘*’ assigned for ‘Yellow Not Pure DCIS’ and ‘Orange IDC’ indicates samples used in the analysis comparing gene expression of DCIS vs IDC for co-occurring patients. ‘Blue Not Pure DCIS’ and ‘Red IDC’ are from tissue biopsies that did not have co-occurring DCIS and IDC in the same sections and were therefor not used for this expression analysis. (**C**) Boxplots illustrating per sample expression data for highly differential genes found when comparing samples in the Early group (E1-E2) with those in the Late group (L1-L2). Differential expression analysis was done using limma-voom and p-values were adjusted for multiple testing using Benjamini-Hochberg correction. Centre line represents the median, box limits represent upper and lower quartiles, whiskers represent 1.5x the interquartile range. Each point represents a sample.

To explore this idea, we used these same 53 selected genes that best separated DCIS from IDC (Adj.P <0.00001) to perform a pseudo-time analysis using a fitted principal curve onto a PCA plot of all our samples (Fig. 3A). We saw that the normal and benign epithelial tissue samples aggregated towards one end of the fitted curve and IDC tissue and DCIS with co-occurring IDC clustered at the other., despite the normal and atypia samples not factoring into the selection of these genes. We then ordered all tissue samples, normal, benign, atypia, DCIS and IDC, by their projection onto the principle curve, creating a pseudo-timeline. We created a heatmap showing expression changes for these genes along the pseudo-timeline, with sample order matching that from the projected principal curve (Fig. 3B). This ‘timeline’ of early breast cancer seemed to reveal how fundamental processes were associated with progression toward invasive disease. Position along the timeline was independent of ER/PR/Her2 status. Moreover, triple negative samples, despite clustering away from other samples on a UMAP when using all genes (Fig, 1A), or even a UPMAP created with just these 53 genes (fig. S4), did not drive the separation on a PCA. This analysis therefore captured the major expression changes shared across most patients, rather than any particular subtype. We observed a gradual loss of expression for genes involved in the epidermis/epithelial development, as we transition from the more normal-like/early-stage DCIS to the later stage DCIS samples and IDC samples. This suggests a progressive breakdown of epithelial architecture, most likely reflecting a loss of integrity in the basal epithelium.

We carried out XCell analysis ^24^ to look for changes in cell type contributions that may occur along this transition, and found further support for epithelial loss with a gradual decline in the enrichment for epithelial cells within each sample when placed in the order of the timeline (fig. S6).

To understand better the changes that occur just within DCIS as they progress closer to the transcriptomic patterns of IDC, we compared DCIS samples from the early part of the time line (Fig. 3b E1-E2) with DCIS samples from the later part of the timeline (Fig. 3b L1-L2) we found a number of smooth muscle related genes were down regulated in the later stages with *TAGLN*,

*CALD1*, *MYL9* and ACTA2 being most significant (Fig. 3c, Table S5). In addition, along with the Collagen genes Col1A2 and Col3A1. *LUM* (Lumican), encoding a small leucine-rich proteoglycan found to be associated with EMT, invasion and metastasis ^25^, and HTRA3 (High-Temperature requirement Factor A3) were found to have the greatest fold change (Fig. 3c). Caldesmon (*CALD1*) has recently been shown to be upregulated in the epithelium of mammary ducts in both mice and humans during lactation ^26^ however its role in the progression of DCIS to IDC has not previously been shown. To visualise the protein distribution in samples representative of different stages of the time line we carried out imaging mass cytometry (IMC) (Fig. 4).

**Fig. 4.**
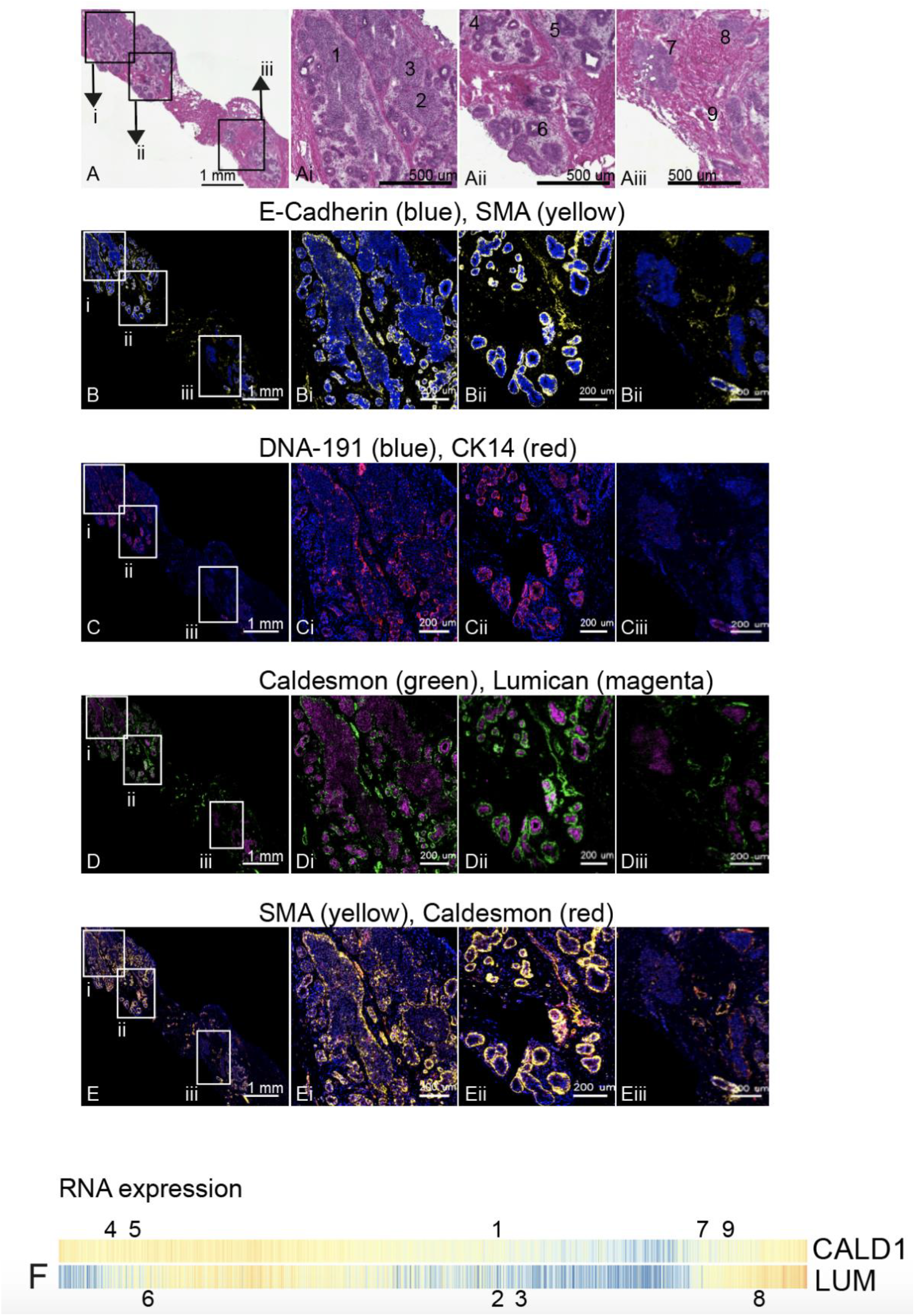
Imaging Mass cytometry of ductal regions from Patient sample CBZ (LumA subtype). The single slide was analysed in one continuous scan and magnified regions retained the same intensity threshold. (A) shows the H&E section, this same section was destained and used for IMC. Boxed areas indicate the corresponding magnified regions shown in Ai,ii and iii. (B) shows E-Cadherin and SMA, with corresponding boxed areas for i, ii and iii. (C) Shows CK14 and nuclear stain DNA-191, with corresponding boxed areas for i, ii and iii. (D) shows Caldesmon and Lumican and (E) shows SMA and Caldesmon, with corresponding boxed areas for i, ii and iii. Numbers on the H&E images indicate regions with expression data from adjacent sections. (F) shows the relative gene expression of *Cald1* and *Lum* (as in Fig.3B) with all samples ordered along the timeline. Numbers above and below pair up with numbers in (A) and mark the position on the Timeline for the two (adjacent) data points corresponding to each region.

As can be seen from the representative slide shown in Figure 4 (where all images are taken from a single slide imaged in one continuous scan), protein staining for Cald1, Smooth Muscle Actin (SMA), and to a lesser extent Cytokeratin 14 (CK14) appear to overlap in localization, and the level of intensity, representative of ion count, is comparative to the relative RNA expression level within the corresponding lesions from the adjacent tissue section (Fig. 4F). Those regions located early in the time line (4, 5, and 6 – parts ii from Fig. 4) have a relatively intact layer of Cald1-expressing cells surrounding the duct, whereas regions further along the timeline (1, 2, 3 and 7 – parts i and iii from Fig. 4) show a much more broken or absent layer of Cald1-expressing cells. Regions towards the very end of the timeline (8 and 9 – parts iii) are starting to upregulate *Cald1* expression and this can be visualised in areas around the duct. The underlying reason for this observation remains unknown however. It is possible that Cald1 is staining fibroblast cells surrounding the ducts, as this marker is frequently associated with this cell type ^27^ . Cald1 positive cells located around regions 8 and 9 (Fig. 4Eiii) may be an alternative form of cancer associated fibroblast (CAF), noted to have a different staining pattern with SMA compared to other regions. This is supported by a recently published study describing a shift in fibroblast phenotype, from normal fibroblasts lining the DCIS ducts, to cancer associated fibroblasts lining DCIS ducts in patients that later developed IDC^9^. A prior study on glioma neovascularization has also described differential expression of splicing variants of *Cald1* in tumour vessels as compared to normal vessels, resulting in upregulation at the protein expression level within the tumour. This was seen to be correlated with a down regulation of the tight junction proteins occludin and Zo-1 – important regulators of mammary epithelia permeability ^28^. As our RNAseq data was not able to reveal transcript variants, we cannot yet attribute this change in expression towards the later end of the Timeline to any particular splice variants. Protein expression for Lumican also appears to follow the trajectory indicated by the timeline, however in this patient, is strongest in the earlier parts of the timeline.

Previous studies have suggested a similar breakdown of myoepithelium during the progression towards IDC, using human breast cancer cell lines and a few select markers ^29^, and very recently a study of human breast tissue with known markers of myoepithelial cells^9,^, lends support to the broader set of expression changes that can be referenced to our timeline.

### The epithelial to mesenchymal transition marks both the early and late stages in the timeline

The timeline revealed a wave in expression of genes relating to the extracellular matrix and cell adhesion, suggestive of a migratory phenotype, initiating relatively early along the continuum and one later, coinciding with the inclusion of the IDC samples (see Fig. 3B). As prior studies ^30^ have indicated that multiple DCIS lesions within an individual patient may be of shared origin, one might imagine that an early loss of adhesion might facilitate spread throughout ductal networks, indeed ∼40% of patients with a DCIS diagnosis, are found to have multifocal disease ^31^, as defined by more than one distinct site of DCIS. Subsequent proliferation and filling of ducts may see a return of cell adhesion with a loss of this property again preceding or coinciding with invasion.

To gain a greater understanding of processes that could be occurring along the timeline, we applied the entire transcriptome, to the MSigDB Hallmarks database to look for gene set signatures. We found the expression pattern of genes associated with the Epithelial to mesenchymal transition (EMT) hallmark signature to closely mirror many of those genes in our time line (Fig. 5A and fig.S8), and also reflected the position of the principal curve along PC2 in our PCA (see Fig. 3a). It has long been proposed that cells within DCIS lesions undergo an EMT along their path toward invasiveness, however, the ability to position our samples along a disease trajectory has allowed us to detect that EMT not only occurs in samples along the timeline at the transition to invasive disease, but also at a second time, much earlier in the disease timeline, when the epithelial architecture surrounding the duct presumably remains intact (region E1 to E2 of Fig. 3B).

**Fig. 5.**
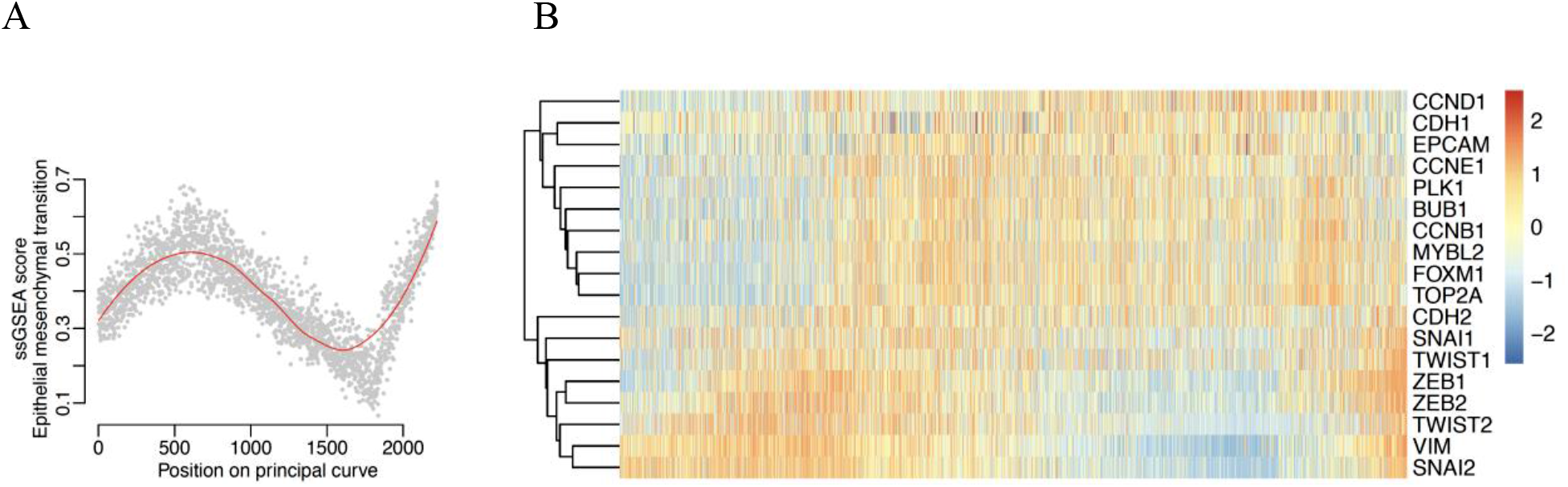
Predominant Hallmark signatures that vary along the timeline. **(A)** Single sample GSEA (ssGSEA) score for the Epithelial to Mesenchymal transition Hallmark signature. Samples are ordered according to the projected principle curve. **(B)** Heatmap showing expression of key proliferation genes (CCND1 – Top2A) and key EMT (CDH2-SNAI2) genes. Samples were ordered according to the projected principal curve. Relative expression is provided as log2 CPM minus the mean log2 CPM for each gene.

**Fig. 6.**
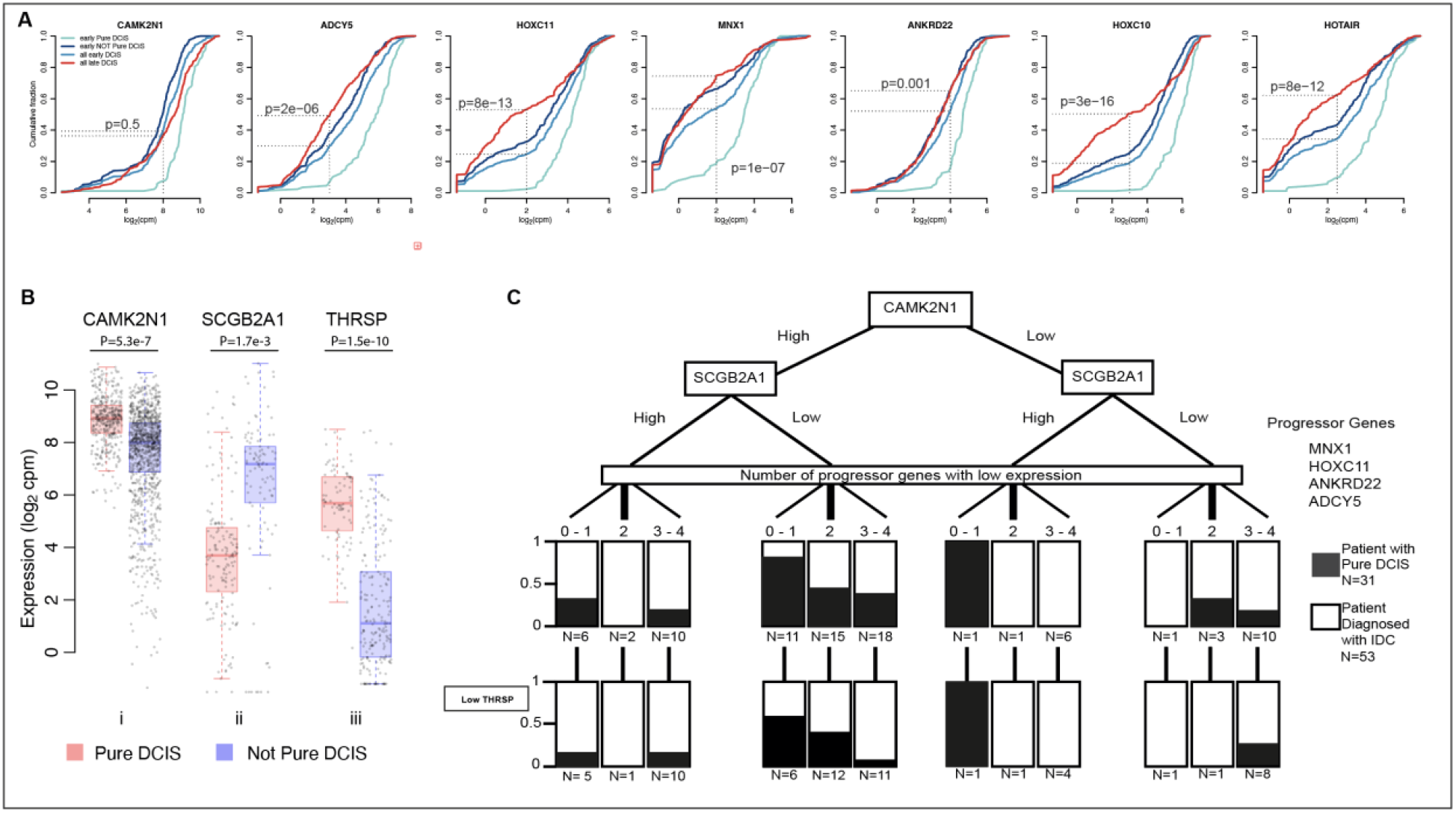
Genes displaying potential as indicators of progression from DCIS to IDC. (A) Cumulative frequency plots for differential genes between early timeline Pure DCIS and early timeline Not Pure DCIS. X axis shows the gene expression in log_2_ counts per million (CPM), Y axis shows the cumulative fraction of samples with the corresponding expression value or lower. Significance values reflect the Fisher’s test for a difference between cumulative fraction of all early DCIS compared to all late DCIS. (**B**) Expression of i. *CAMK2N1* for all DCIS samples, ii. of *SCGB2A1* for all patients in the Low Hazard group – 1 progressor gene down regulated and *CAMK2N1* high, and iii. *THRSP* for all patients 3-4 progressor genes down regulated, *CAMK2N1* high and *SCGB2A1* low. Centre line represents the median, box limits represent upper and lower quartiles, whiskers represent 1.5x the interquartile range. Each point represents a sample. (**C**) Separation of patients with no IDC identified in our tissue sample. 31 patients were never diagnosed with IDC after 10+ years, 53 patients were diagnosed with IDC in a secondary biopsy. Black/ white regions reflect the proportion of patients with each diagnosis (Pure DCIS vs with IDC) within each node. Boxes in the low *THRSP* layer reflect the number of THRSP low patients from the node above.

The emergence of EMT at very early time points in disease could suggest that cells require this process to migrate though the ductal system, disseminating and forming multifocal DCIS. Following an early dissemination phase, cells may again adopt a more epithelial character as they become proliferative, with a later acquisition of mesenchymal features coinciding with exit of tumour cells from the duct. The possibility of both an early and late EMT phase could be something to consider when using EMT markers to group DCIS cells into those that may be pre-invasive versus more indolent. EMT potentially occurring twice during the progression from normal epithelium to IDC might suggest that it is alone insufficient to enable invasion but that it must be coupled to breakdown of the myoepithelium for transformed cells to escape from the confines of the duct.

### Cell proliferation increases after the early EMT phase

We also identified additional processes within the MSigDB Hallmarks database that were not represented by our 53 Timeline genes, yet still correlated with changes in tissue states along the path between normal epithelium and invasive disease. We observed what appeared to be an altered regulation of the G2/M checkpoint signature in the early stages of the timeline (fig. S7), however only a subset of genes were actually contributing to the signal. On closer examination we found that these genes were all associated with proliferation, including genes identified as being key to the proliferation signature (*MYBL2*, *BUB1* and *PLK1*) ^32^. This increase in expression of proliferation genes appears to initiate just after the first peak in expression of EMT related genes (Fig 5B), supporting the notion that after migration through the ducts, cells resettle proliferate as they occupy new sites.

### Reduced expression of *GLTSCR2* and perturbation of ribosomal biogenesis is an early DCIS event

As we generated what appeared to be a progression timeline, we next sought to identify genes that may be altered during the earliest stages of disease initiation. For this we focused first on the DEGs between all normal (hereafter normal refers to the non-neoplastic normal and benign tissues) tissue samples and all Pure DCIS (with the notion that Pure DCIS samples are less likely to be influenced by the transcriptional changes that come with the presence of IDC) . We then looked for shared genes also significant between normal and DCIS only using samples in the very early part of the timeline (prior to E1 in Fig. 3). In doing this we retained the added strength of a large data set by using all samples, but removed the strong expression signature that arose from the onset of increased proliferation and EMT (that came after E1 in Fig. 3) . We found *GLTSCR2*, also known as *PICT-1*, to be the most significant DEG when using all normal and all Pure DCIS samples (FC; 1.7 Adj. P; 2.8e-69) and more highly expressed in the normal tissue samples (fig. S9). This was also one of most significant DEGs in the very early timeline samples (FC; 0.9, Adj. P; 1.6e-14). GLTSCR2, is thought to act as a tumour suppressor ^33,34^ and has been shown to translocate to the nucleoplasm, provoked by ribosomal stress, where it interacts with, and stabilizes, p53 to inhibit cell cycle progression ^35^. Decreased expression was seen to delay DNA repair and abolish G2/M checkpoint activation ^33^. The ribosomal proteins *RPL5* and *RPS6* are, after GLTSCR2, the most significantly down regulated genes when comparing all Pure DCIS samples with all normal ductal tissue, (FC; 1.3e-66 and 1.1e-57 respectively), and both genes were also among the most significant DEGs when comparing samples from the very early timeline. In addition to their role in the ribosome, *RPL5* and *RPS6* have been shown to be essential for the activation of p53 in response to DNA damage ^36^. Pairing the top 100 DEGs between all Pure DCIS and all normal samples, with highly significant DEGs (Adj. P < 1e-10) from the same comparison using only the very early samples, we found 44 overlapping genes, with 19 of these related to ribosomal biogenesis (Table S1). Although ribosomal proteins appear to function in a variety of different ways, there is increasing evidence for their role in tumour development ^37,38^, and it is possible that what we are observing at the early stages of the timeline could reflect their involvement in the initiation of DCIS. In addition to ribosomal-related genes, we also observed a significant down regulation of the transcription factor *NFIB,* encoding the Nuclear Factor I B, in DCIS samples, with this gene being the most significant DEG when comparing DCIS with normal epithelium samples taken from the very early timeline (FC; 1.3e-28) (fig. S9). *NFIB* is part of the NFI gene complex, together with *NFIA*, *NFIC* and *NFIX* and recent work has described *NFIC* as being a regulator of ribosomal genes within the pancreas ^39^. However, the ribosomal genes affected by *NFIC* share very little in common with the genes we find most differential in our analysis, and as yet no other work has associated *NFIB* with modified expression of ribosomal genes, thus the expression changes here could be reflective of an additional process in early disease.

Current understanding of the transcription factor, Nfib, in breast cancer associates the over expression of this gene with metastasis ^40,41^, however, it has also been demonstrated using a prostatic mouse model that heterozygous and homozygous loss of Nfib can lead to epithelial hyperplasia ^42^. RNAseq analysis from this same study, comparing *NFIB*^-/-^ to *NFIB*^+/+^ prostatic grafts, identified 138 DEGs, some of which, such as *FOXC1* and *SOX10,* are also differential in our analyses of both early timeline samples, and in all normal vs DCIS samples, suggesting a shared role for *Nfib* in both prostate and mammary epithelial tissue.

### Progression along the disease timeline follows divergent paths depending on hormonal status

In contrast to the early stages of disease, we see a divergence in dominant hallmark signatures later in the Timeline when we look at samples grouped by oestrogen receptor status (fig. S7). Not surprising the Oestrogen Response signatures are up in ER+ samples as they progress closer to IDC, and this is not observed in ER-samples. The later stage of the timeline for ER-samples appears to engage an immune response as reflected by a substantial rise in both the Interferon Gamma and Interferon Alpha response signatures. We also see a reduction in the Oxidative Phosphorylation signature in ER-samples.

### Potential indicators of progression competence within early-stage lesions

The ability to discriminate DCIS lesions that have a higher potential to progress to invasive disease would have enormous impact in the clinic. We therefore asked whether we could identify indicators of progression potential that could be used even if a patient presented with DCIS and no evidence of invasive disease. The position of a patient’s DCIS sample along the timeline did not appear to be indicative of that patients’ diagnosis, i.e. Pure DCIS or IDC (mean difference in position on the timeline between Pure DCIS and Not Pure DCIS – 130; p-0.11; Welch two sample t-test), this is in contrast to a recent study, describing Pure DCIS patients as having a less intact myoepithelium as compared to those that later developed IDC^9^, indeed we did not see an enrichment for Pure DCIS patients in the later end of our timeline where samples display reduced expression for epithelial related genes ^43^. Having the transcriptomes of micro-dissected lesions ordered along a timeline of progression however, offers the opportunity to probe a more comprehensive dataset with unbiased markers. Given that our timeline indicates a distribution of DCIS expression phenotypes, we examined DCIS samples from three groups: those ‘early’ in the timeline (region E1 to E2, Fig. 3B), in the middle of the timeline (between E2 and L1) and late in the timeline, adjacent to the IDC-enriched region (region L1 to L2). Comparing the transcriptome of Pure DCIS to Not Pure DCIS revealed 308 DEGs for samples within the early part of the timeline, 206 for the mid region, and just 90 for the late stage of the timeline. The difference in the number of DEGs as we progress along the Timeline supports our ordering of samples and suggests that the distinction between samples derived from Pure DCIS patients and patients where DCIS is associated with invasive disease becomes less apparent as the disease progresses along the timeline. This might be expected if lesions are converging on a phenotype similar to that of invasive disease. Interestingly, comparing DCIS samples with IDC samples, both from the very late region of the Timeline (past L2) found no consistent DEGs.

To search for potential markers that could distinguish patients who would be more or less likely to progress to IDC, we first looked at the early region of the timeline, comparing samples from those patients with Pure DCIS to those patients who were diagnosed with IDC (concurrent, or at any timepoint after the biopsy was taken) – NOT Pure DCIS . We identified DEGs where the DCIS samples associated with an IDC diagnosis had a bimodal or skewed distribution of expression values, and the samples from Pure DCIS patients had an oppositely skewed pattern. We identified 7 such genes: *CAMK2N1*, *MNX1*, *HOXC10*, *HOXC11*, *ADCY5*, *ANKRD22*, and *HOTAIR*. All showed a distribution of expression values that were lower in the DCIS associated with IDC samples as compared to Pure DCIS (Fig. 6A). If these genes were early indicators of progression potential, one might imagine that their expression changes would be enriched among all DCIS samples as they became more similar to IDC along the timeline. We therefore compared the distribution of expression values in all DCIS samples from the early part of the timeline (region E1 to E2, Fig. 3B) to all DCIS samples from late in the timeline (region L1 to L2). To differing degrees, all except *CAMK2N1* showed a general decrease in the distribution of expression values in later stage samples (as defined by the timeline, Fig. 6A).

Differences in the distribution of expression values for *CAMK2N1* were exclusively linked to patient status (Pure DCIS versus Not Pure DCIS). Its expression remained discriminatory in all stages of the timeline, though it did not reach significance in later stages. This gene encodes a recently identified inhibitor of Calcium/calmodulin-stimulated protein kinase II, a protein thought to be involved in various cellular processes including cell proliferation, mammary gland lumen formation, and cancer cell metastasis. Expression of this protein kinase (*CAMK2*) is also predictive of poor breast cancer patient prognosis ^44^. *CAMK2N1* itself has been reported as a prognostic marker for ovarian cancer ^45^ and plays a tumour suppressive role in prostate cancer ^46^ and glioma ^47^, and in comparing all Pure DCIS with all other DCIS samples, is significantly down regulated in Not Pure DCIS samples (Fig. 6B).

*HOXC11*, *HOXC10* and *MNX1* each contain a homeobox domain, and *HOTAIR* is an antisense RNA whose source locus is found within a cluster of *HOXC* genes, between *HOXC11* and *HOXC12*. Homeodomain proteins function as transcription factors, regulating gene expression and cell differentiation during development, and have been frequently associated with cancer progression, where they are either up or down regulated, depending on the Hox family member and cancer type. A recent study modelling the growth expansion of DCIS posited an initial rapid expansion phase, followed by a long-term steady phase were cells were predicted to be in a cell density induced quiescent state ^48^. Notably, down regulation of *HOXC10*, *HOXC11* or *MNX1* has been reported to reduce cell proliferation in a variety of different cancers ^49-52^, so could suggest a possible quiescent state prior to invasion. Similarly, knockdown of the ankyrin repeat domain 22 gene, *ANKRD22*, inhibited the proliferation, invasion and epithelial-to-mesenchymal transition of breast cancer cells ^53^, and a number of studies have reported high levels of expression being associated with poor outcome in non-small cell lung cancer ^54^ and prostate cancer ^55^, an inverse correlation to what we observe here with a ductal in situ disease. The adenylate cyclase 5 gene, *ADCY5*, is thought to be regulated by the tumour suppressor gene *FOXP1*, and knockdown of *FOXP1* was followed by a significant upregulation of genes attributed to chemokine signalling pathways, including *ADCY5* ^56^.

*HOTAIR* has previous been identified as a segregation marker between two clusters of DCIS ^57^, however this prior study noted that an upregulation of *HOTAIR* was associated with a more ‘aggressive’ cluster of DCIS. This aggressive cluster however, was predominantly triple-negative disease, whereas our groups were not segregated by subtype, and the DCIS samples in the latter part of our timeline were predominantly not triple negative. Other studies have reported an upregulation of *HOTAIR* when comparing human cancers to adjacent non-cancerous tissue ^58^, and we also found that this LncRNA showed lower expression in our normal epithelium samples, albeit at levels similar to what we see in the DCIS associated with IDC samples from the early part of the timeline. .

As *HOXC10*, *HOXC11* and *HOTAIR* loci are closely linked on the same chromosome, it seemed possible that the changes in expression that we observe could have resulted from copy number loss; however, we do not see a similarly reduced expression for *HOXC12* or *HOXC8*, the two adjacent genes.

To provide a foundation for future validation studies, we wondered whether we could use any combination of these markers to associate patients from this study, with the presence of IDC. We formulated a decision tree, focusing on protein coding genes which may be more routinely evaluable clinically. Because of its ability to segregate the samples from the Pure DCIS group from the Not Pure DCIS group in all timeline categories, we placed *CAMK2N1* at the top of the tree, separating high and low expression categories. We then explored different ways of using information on the expression of *MNX1*, *HOXC11*, *ANKRD22*, and *ADCY5*, as in no scenario did *HOXC10* seem to add additional discriminatory power. We found that simply tallying the number of these ‘progressor’ genes that were down-regulated enabled us to bin patients into groups within the decision tree. These 4 markers, plus *CAMK2N1*, enriched for patients who did not progress to IDC by 3.6-fold using the criteria of 0 - 1 gene down regulated and *CAMK2N1* high (Lower Hazard group) (36% vs 10%, - patients from the Pure DCIS group vs patients with an IDC diagnosis), whereas 3-4 genes down regulated or *CAMK2N1* low (Higher Hazard group) enriched for patients that received a diagnosis of IDC by 1.7-Fold (71% vs 42% - patients with an IDC diagnosis vs patients from the Pure DCIS group) (fig. S10). This difference within the Higher Hazard category, may suggest that many more patients might have progressed to IDC had they not been treated. This percentage is consistent with current research suggesting that between 13-53% of patients with untreated DCIS will progress to invasive disease ^59^. Interestingly, we noticed that the majority of Her2-positive patients in the Pure DCIS group fell into the Low Hazard group (6 out of 8); however, we did not see this enrichment in the Not Pure DCIS group. Previous studies have noted a higher proportion of Her2-positive DCIS cases compared with that seen in invasive disease, and it has previously been suggested that a Her2 DCIS may actually be less likely to progress to IDC ^60^, our findings here would support this hypothesis. We next sought to identify additional markers that could segregate the Lower Hazard group, further differentiating those patients with Pure DCIS from those diagnosed with IDC. We found *SCGB2A1*, encoding Mammaglobin B, to be significantly differential between the two groups and able to provide further discrimination between patients (Fig. 6B). High expression of *SCGB2A1* was frequently associated with high expression of *SCGB2A2* and *SCGB1D2*, encoding Mammaglobin A and lipophilin B. Expression differences at both the RNA and protein level of SCGB1D2 have also been observed in a prior study of 24 patients, comparing DCIS with and without progression to IDC^5^. All three proteins are members of the secretoglobin superfamily and are known to be upregulated in breast cancer, with SCGB2A2 and SCGB1D2 forming a multiprotein complex ^61^. Using this additional marker, we were able to place 29% of Pure DCIS patients into the Lower Hazard group whereas just 3% of those with IDC fell into the Lower Hazard group (Table S2 shows expression values for high and low expression of each gene). Taking the subset of patients, where we found only DCIS in our tissue biopsy (DCIS with IDC patients and Pure DCIS patients), and blinded by any diagnosis of IDC from other tissue biopsies from the same patient, we were also able to discriminate those who had been diagnosed with IDC using our markers (Fig. 6C).

We next sought to understand why some patients with Pure DCIS, while being grouped into the Higher Hazard category, according to our marker set, had however, not been diagnosed with IDC. For this we first compared all patients high for *CAMK2N1* and low for *SCGB2A1*, with reduced expression of 3-4 progressor genes (N = 25 patients diagnosed with IDC; N = 7 patients with Pure DCIS). We found *PHGR1*, *THRSP* and *SERPINA5* to be highly differential between the two groups, with increased expression in Pure DCIS (Fig. 6B and Table S3). Although these genes were frequently co-expressed, we found *THRSP* able to best segregate the Pure DCIS patients from those patients diagnosed with IDC (Fig. 6C). We did not find this gene to be additionally informative for any other group on the decision tree. *THRSP* encodes the Spot14 (S14) protein, which regulates fatty acid synthesis in mammary epithelial cells ^62^. Over expression of this protein was seen to reduce the tumour latency period in mice and increase proliferation; however, this same study showed an overwhelming reduction in lung metastasis in these same mice compared to controls or *THRSP* knockout mice. This gene, along with other genes involved in fatty acid biosynthesis, was also found to be down regulated in invasive growth compared to in situ growth in a mouse model of DCIS^63^. Similarly, upregulation of *SerpinA5* has been linked to reduced metastatic and invasion potential in both ovarian and breast cancer ^64,65^. Comparing Pure DCIS with Not Pure DCIS for all samples with reduced expression for 3-4 progressor genes we found a number of DEGs (Table S3) previously associated with invasion and metastatic potential that were expressed at consistent levels (correlating with reduced metastasis) for all Pure DCIS samples, including *SERPINE2* and *SLPI*, both genes found to influence metastasis and contribute to vascular mimicry in a mouse model of breast cancer ^66^. These Pure DCIS samples were also predominantly located in the later stage of our timeline (L1-L2 region), suggesting they may lie at the point when they need to acquire the capacity to leave the duct as the next step in their progression.

The functional diversity of these markers may indicate that multiple factors must come together for DCIS to progress to invasive breast cancer. Although we have proposed possible progression markers that will require more extensive validation, it still remains to be seen whether, and how, each of these may play a role in this disease. A recent study ^7^ also looking at potential biomarkers of DCIS progression, identified the genes *FGF2*, *GAS1* and *SFRP1* as being markers of in situ progression, suggesting their downregulation contributed to the invasiveness of epithelial cells. In support of this we also see that these 3 genes are notably down regulated as samples are arrayed along the timeline. These previously described genes, although discriminatory between DCIS at early, and DCIS at later points on the timeline, were not differential between Pure DCIS patients and DCIS patients diagnosed with IDC, at any time point on the timeline.

## Discussion

Though widespread screening for breast cancer has detected disease in many more women at an early stage, a corresponding decrease in breast cancer deaths has not been forthcoming. (Breast Cancer Facts and Figures 2019-2020 – American Cancer Society). Instead, many more women are receiving treatment for non-invasive disease, which may include chemo- or radiotherapy, coupled with breast-conserving surgery or mastectomy. Numerous studies indicate that a substantial fraction of women with a diagnosis of DCIS would never progress to life-threatening invasive disease^67,68^. Therefore, many women are being needlessly overtreated using therapies with significant and long-term deleterious side effects. This realization provokes an urgent call for a better understanding for the development of DCIS and ways to discriminate those who will progress to invasive disease, and thus require more aggressive treatment, from those who are unlikely to do so and who may opt for less extensive interventions. Our transcriptomic analysis of this large data set has enabled us to identify processes that may characterize the progression of DCIS from initiation to invasive disease and to identify candidate biomarkers, which may be associated with progression. Though these particular markers will need to be validated in independent and larger cohorts, the generation of these hypotheses illustrates the utility of this large-scale dataset for the broader community.

## Methods

**Extended material and methods information is available within the relevant sections within supplementary information.**

### Material collection

Freshly frozen tissues were donated for research by a cohort of women having undergone a medically indicated diagnostic breast core biopsy, following an abnormal mammogram with suspected malignancy. Multiple adjacent sections were cut from each tissue core. Guided by pathological annotation, regions of IDC, DCIS, atypia, benign, and normal epithelium were isolated by laser-capture micro-dissection with regions of the same individual lesions taken from 3 adjacent sections. RNAseq libraries were made using the SMARTer ultra-low RNA kit V3 (Takara Bio USA, Mountain View, CA, USA). and analysed individually (i.e., lesions from adjacent sections were not pooled) from each sample region and were quality filtered. A total of 2222 libraries from 143 patients passed our quality metrics and were taken forward for subsequent analyses (Table S4). Each sample was classified into the generally accepted subtype groups (Luminal A, Luminal B, Basal-like, Her2-enriched and Normal-like) using Absolute Intrinsic Molecular Subtyping (AIMS) (fig. S1) ^10^.

### Patient and sample group assignment

Patients were assigned to one of four categories; Pure DCIS - where ipsilateral IDC had not been reported in the patient, neither at the time, nor in follow up appointments during more than 10 years since original diagnosis (details Table S4), N = 31; DCIS+IDC - Where the biopsy dissected featured both DCIS and IDC lesions, N=45; DCIS with IDC - Where the biopsy dissected only featured DCIS however the patient had been diagnosed with IDC (from clinical or pathology biopsies, or at a later time), N = 55; IDC – where no DCIS was found in the dissected biopsy, but had been diagnosed in additional biopsies; N = 2. Or where no DCIS was diagnosed in any of the biopsies N = 4. normal epithelium, benign ducts, and atypia were taken from the same biopsies as above where present in the section or from additional patients diagnosed with DCIS in other biopsies (fig. S2 shows clustering of all samples). Samples coming from patients in categories DCIS+IDC and DCIS with IDC are collectively grouped as ‘NOT Pure’.

### Differential expression analysis

Differential expression analysis was done using Limma-Voom, due to its ability to handle large datasets and replicate measurements for the same sample. First, expressed genes were selected by the ‘filterByExpr’ function. Calculation of normalization factors was carried out using the TMM method. To correct for multiple samples coming from the same patient, we used a double ‘voom’ approach, including a ‘duplicationCorrection’ step with blocking based on patient. Fitting was done using ‘ImFit’ (with blocking and correction applied if applicable), followed by construction and calculation of contrasts using ‘contrast.fit’ function followed by ‘eBayes’. A gene was considered to be differentially expressed if the Benjamini-Hochberg adjusted p-value was <0.05.

### Pseudo-time analysis

A differential expression analysis was carried out as described, between DCIS and IDC samples, taken from only those patients with data from both tissue types. This was followed by a Principle Component Analysis (PCA) using only the most significant genes (p<0.00001, n=53). Remaining samples for DCIS, IDC, normal, benign and atypical epithelium, were then embedded onto this PCA to reveal the major patterns in the data and avoid grouping solely based on patient. Since the different tissue types were positioned on the PCA in a biologically meaningful order, we fitted a principle curve to the data and projected the samples onto the curve to arrange them by their predicted pseudo-time order. Linear methods such as ordering along PC1 have previously worked well for single cell data^69^ and we did not expect to be able to describe bi- or multifurcation events. We chose a principal curve over using PC1 to order the samples due to its ability to capture the expression wave along PC2. We note that arranging the samples according to their position on a UMAP embedding resulted in largely the same order (see fig.S5).

### Gene set enrichment analysis

The R Bioconductor package RITAN (v.1.10.0) was used for gene set enrichment analysis using the MSigDB Hallmarks database. All protein-coding genes were used as a background. Terms with FDR-adjusted p-value < 1e-5 are listed. To determine enrichment across the timeline, we used a sliding window of 100 samples, moving 50 samples at a time, compared to all remaining samples. For the ssGSEA we used the GSVA package^70^ from R Bioconductor with the “ssgsea” method.

### Imaging Mass Cytometry (IMC)

Previously stained H&E slides (used for annotation purposes prior to LCM) were first de-stained using a combination of ethanol and acidic ethanol (see Supplementary methods, IMC section for further details). Heat induced antigen retrieval was carried out in Tris-EDTA for 20 minutes, slides were then cooled and blocked prior to overnight incubation with metal conjugated antibodies;

Anti-Caldesmon (Abcam; ab215275), Anti-Lumican (Abcam; ab198974) Anti-cytokeratine 14 (ab236439), Smooth Muscle Actin (SMA) (Thermofisher; 14-9760-82) and E-Cadherin (BD Biosciences; 610182). Slides were then imaged using the Hyperion Imaging Mass Cytometer (Fluidigm). IMC datasets, saved by the Hyperion instrument as .mcd files, were initially converted to the *Zarr*format, preserving the entire signal dynamic range and metadata, using a custom python script described previously^71^ and available at https://github.com/IMAXT/imc-nuclear-segmentation. The resulting zarr datasets were visualized using a custom-made IMC viewer tool, also written in python, operating on a jupyter notebook instance (also described in the same publication and available at https://github.com/IMAXT/imaxt-image).

### Patient marker classifier for group assignment on the decision tree

High and low expression of each marker gene was based on the majority segregation between Pure DCIS and Not Pure DCIS. Table S2 provides additional information regarding the expression levels for each gene. A patient was placed in a group based on a minimum of 2 samples representing the ‘associated with IDC” expression levels, this being low *MNX1*, low *HOXC11*, low *ANKRD22*, low *ADCY5*, High *SCGB2A1*, low *Camk2N1* and low *THRSP*. Two patients were removed from the decision trees as data was only available for 1 sample.

## Acknowledgements

We are grateful for the support provided by the project management team at the New York Genome Centre, namely Maddalena Coppo, Elisa Venturini, Sam Phillips, and Catherine Reeves. We thank Laurence de Torrente, Bassem Ben Cheikh, and Simon Knott for their contributions to the project as a whole, and Dario Bressan and Fatime Qosaj for their guidance and contributions to the IMC part of the project. We would also like to thank the histopathology/ISH core facility and the Genomics core facility at Cancer Research UK, Cambridge Institute, for their support throughout the entire project, and the team at Cedars-Sinai, Los Angeles, namely Steven Piatidosi and Andre Rogatko, for their role in the clinical databases.

## Authors contributions

Conceptualization: CR, GH, HKL

Data curation: CR, SB, HG, AH, NR

Formal analysis: SB

Funding acquisition: CR, JG, GH, HKL

Investigation: CR, JX, JG, JFA, ER, AN

Projection administration: CR, AH Resources: NR, AH, HKL

Supervision: CR, GH Visualization: CR, SB

Writing – original draft: CR, SB

Writing – review & editing: CR, JX, SB, JG, NR, HKL, GH

## Data availability

Raw sequencing data (aligned to GRch38/hg38) that support the findings of this study have been deposited in European Genome-Phenome Archive under the study ID number EGAS00001005370, and are available on application to the Data Access Committee upon request to. Most additional data supporting the findings of this study are available within the paper and its supplementary information files. The remaining data are available from the corresponding author upon reasonable request.

## Supplementary Materials

## Supplementary Materials and Methods

### Patient tissues

Freshly frozen breast tissue was analysed under a Duke University IRB approved Tissue Use Protocol Pro00059726. These biopsies were originally consented for tissue banking and study under the Duke Breast SPORE grant (Pro00014678), the DUHS Biospecimen Repository and Processing Core (BRPC) Facility protocol, the DOD TVA tissue bank (Protocol #Pro00045965), or the DOD CTRA tissue bank (Protocol #Pro00044981). Primary breast cancer specimens were collected from women with an abnormal mammogram suspicious for malignancy and undergoing a medically indicated diagnostic breast core biopsy sampling who were willing to donate cores of tissue for research. After obtaining informed consent, a diagnostic core biopsy was conducted, and additional research cores were obtained. The research cores were frozen immediately in OCT embedding compound in the vapor phase of a liquid nitrogen bath or on dry ice and held frozen at -80 C until a definitive diagnosis was made by pathologic assessment of the diagnostic cores. At time of definitive diagnosis, H&E stained frozen section slides were prepared from the research core biopsies and compared with the results from the diagnostic cores by a pathologist with expertise in breast pathology. Tissue was stored in a locked and monitored -80C freezer until it was used for this study.

### Tissue preparation

Frozen tissue biopsies were sectioned under RNAse clean conditions. Ten serial sections of each were taken, with two sections per slide – 6 sections (10µM) on PEN slides and 4 sections (5µM) on glass slides. The first and last (glass) slides were subjected to H&E staining, mounted and annotated by an experienced pathologist. Remaining sections were mounted on PEN slides, and stored for a maximum 1 week, before H&E staining immediately prior to micro-dissection.

### H&E staining

Sections were fixed in 75% ethanol for 40 seconds followed by 30 seconds in RNAse free water. Sections were then treated with Hematoxylin solution (Harris Modified, Sigma-Aldrich) for 30 seconds, washed in water for 30 seconds in three different containers, before being dipped into Blueing reagent (0.1 % NH_4_OH, Sigma-Aldrich ) for 30 seconds followed by Eosin solution (Sigma-Aldrich ) for 10 seconds. Lastly sections were dehydrated in rising ethanol concentrations (70, 95 and 99.5% ethanol, 30 seconds each) and air dried.

### Laser capture micro-dissection

Lesions were first paired up with the pathologist annotated regions, and each lesion was identified in all tissue sections prior to dissection. IDC lesions (and occasionally DCIS lesions) were more variable in their distribution through the sections and no lesion was dissected if it was not clear that we could identify the same lesion in the neighbouring section. Tissues were cut using a drop in the tube cap-laser dissection (LCM) microscope (Leica DM6000R/CTR6500) using the Leica LMD7000 system (Leica Microsystems CMS GmbH, Wetzlar, Germany). Images were taken (and confirmed by the pathologist) and cells were dissected under 10X or 20X magnification, with the minimal laser power necessary. Isolated cells were collected in 9µl of lysis buffer (for RNAseq library preparation). The tubes were then snap frozen on dry ice (with tissue remaining in the cap) and tubes stored upside at - 80 °C until further processing. Lesions were collected over 3 adjacent sections and each individual dissection corresponded to 1 RNAseq library preparation, for example a biopsy with 3 DCIS containing ducts had 9 individually dissected regions, 9 RNAseq preparations and represented 9 samples for expression data, which were then subject to the below described quality filtering.

### Imaging Mass Cytometry (IMC)

#### De-staining of H&E slides

H&E-stained slides were submerged in 100% xylene for up to 72 hours (less time for those slides more recently stained with H&E), to ease the removal of the coverslip. Slides were rinsed twice, one minute each in fresh 100% xylene to remove any residual adhesive. Following, slides were rehydrated in fresh absolute ethanol, 3 times for 1 minute each, to remove the eosin stain. Slides were then placed in 1% acid alcohol (HCl in 70% ethanol) for one minute with gentle agitation to remove Haematoxylin stain. Finally, slides were rinsed twice in water.

#### Tissue antibody labelling

De-colourised H&E-stained sections (5uM) initially used for tissue annotation before the laser capture microdissection in the study were re-used for antibody labelling. The slides were directly subjected to antigen retrieval after H&E de-staining. Heat-induced epitope retrieval was conducted in a water bath at 95°C in Tris-EDTA (pH=9.2) buffer for 20 minutes. Following cooling, slides were blocked with 3% BSA (Sigma) in TBS containing 0.3% Triton X-100 (Sigma-Aldrich) for 1h at RT. Slides were then incubated with metal-tagged antibodies overnight at 4°C. Following incubation, slides were first washed twice with TBS/0.1%Tween 20 and then twice with TBS. Finally, slides were rinsed once with water and incubated with 0.5μM Cell-ID Intercalator-Ir (Fluidigm, 201192B) at RT. After 15 minutes slides were briefly rinsed with water and air-dried for at least 30 minutes before IMC acquisition.

#### Antibodies

Lanthanide metal-labelled antibodies conjugated in-house using Fluidigm’s MaxPar’s antibody conjugation kit (Fludigm).

Anti-Caldesmon (Abcam ab215275) - this antibody has been validated by abcam, including with a knockout validation, it also stains positive regions that correlate with other publications (e.g. Stevenson et al. 2020 in references) and the Human Protein Atlas which use different Caldesmon antibodies.

Anti-Lumican (Abcam ab198974) - This antibody has been validated by Abcam with WB and Flow. There is no direct validation in our tissue of study, however, the intensity level does partly correlate with RNA expression level from adjacent sections.

Anti-cytokeratine 14 (CK14) (Abcam ab236439) - this antibody has been validated by abcam, including with a knockout validation, it also stains positive regions that correlate with multiple publications and the Human Protein Atlas , all of which use different ck14 antibodies.

Smooth Muscle Actin (SMA) (Thermofisher 14-9760-82) - This antibody has been validated by Thermo and others that have submitted publications using this antibody (32). There is relative expression and WB validation. It also stains positive regions of the breast that correlate with multiple publications and the Human Protein Atlas , all of which use different SMA antibodies.

E-Cadherin (BD Biosciences 610182) - This antibody is noted by BD to have some degree of cross reactivity to P-Cadherin. With both proteins sharing localisation within the mammary gland as shown by The Human Atlas. The use of this antibody is more for visualization of the tumour location, and we are making no reference to the level of staining for this antibody.

#### Imaging mass cytometry acquisition

IMC images were acquired using the Hyperion Imaging Mass Cytometer (Fluidigm). The air-dried slide was loaded into the imaging module, where an optical preview of the ROIs was recorded for laser ablation. Tissues were ablated by a UV-laser spot-by-spot, line-by-line at a resolution of 1 um and a frequency of 200 Hz. Further details can be found within the main methods sections.

### RNA sequencing Preparation

Samples were processed according to manufacturer’s instructions with 15 cycles of pcr amplification using the SMARTer ultra-low RNA kit V3 (Takara Bio USA, Mountain View, CA, USA). Amplified cDNA was fragmented using the Covaris LE220 sonicator (Covaris, Woburn, MA) according to the manufacturer’s instruction to yield a target fragment size of 200 bps. The sequencing library was then prepared from fragmented cDNA using NuGEN Ovation Ultralow Multiplex System (NuGEN, San Carlos, CA, USA) with 12 cycles of PCR. Finished libraries were purified from free adaptor product using RNAClean XP beads (Beckman Coulter Genomics, Brea, CA, USA). The resulting purified libraries were quantitated using a Qubit (Thermo Fisher Scientific, Waltham, MA USA) and the Kapa library quantification kits (Roche Life Science, Indianapolis, IN USA). The size range of the libraries was confirmed by the Agilent 2100 Bioanalyzer and the Agilent 4200 TapeStation (Agilent Technologies, Palo Alto, CA, USA). An equal amount of DNA was used to pool up to 6 samples per pool.

### RNA-seq alignment and quantification

Raw reads were aligned to the GRCh38/hg38 reference genome using STAR (*65*), (v2.5.2, --alignIntronMax 200000 --alignMatesGapMax 200000 --chimSegmentMin 15 --chimJunctionOverhangMin 15, Gencode V25 gene models). Gene counts were derived using featureCounts (v1.4.3) with default options and Illumina iGenomes Refseq annotations (corresponding to GCF_000001405.30).

### Quality assessment of RNA-seq data

We obtained 2724 initial samples for analysis after excluding failed libraries with <1 million raw reads, <15 % uniquely mapping reads, or <5 % of the raw reads mapping to genes.

For each tissue type (DCIS, IDC, normal epithelium, benign epithelium, atypical epithelium,) we then applied the following additional filtering. Limma-Voom (*66*) was used to calculate TMM normalization factors and convert the normalized counts to log_2_ counts per million (CPM) values. A three-step filtering procedure was employed to remove low-quality samples based on their global gene expression patterns. First, the Pearson correlation between each sample and the mean log_2_(CPM) was calculated and the worst sample was iteratively removed and the mean re- calculated, this was repeated until all remaining samples had correlation ≥0.70 (≥0.65 for IDC samples due to their increased heterogeneity) to the mean. Second, individual samples that were more correlated to the mean of all samples than to the mean derived from the patient in question were excluded. Third, the correlation between each sample and the mean log_2_(CPM) for samples from same patient, was calculated and the worst sample was iteratively removed and the mean re-calculated until all remaining samples had Pearson correlation ≥0.80 (≥0.75 for IDC samples).

The thresholds were chosen to remove only samples that were either failed or were of considerably less quality compared with other samples from the same tissue and/or patient. In addition, during further validation we noticed that this filtering procedure excluded more basal samples than any other molecular subtype and we therefore opted to use more lenient thresholds for those samples (DCIS samples predicted to be basal were filtered using the IDC thresholds, and IDC samples predicted to be basal were filtered using ≥0.60 and ≥0.70 as thresholds).

In total, 414 samples were removed by the first filter, 43 by the second, and 45 by the third filter, resulting in 2222 retained samples in the final dataset, representing 1230 distinct lesions from 143 patients, with 274 lesions present as a single sample, 902 lesions present as two samples derived from different sections, and 48 lesions present as three separate samples. All samples and filtering results are listed in Supplementary Table 4.

### Molecular subtype classification

Molecular subtypes (Her2, Normal, Basal, LumA, LumB) were assigned using the AIMS package^68^ from R Bioconductor applied on the expression counts matrix. RNA expression levels for *ESR1*, *PGR* and *ERBB2* were established based on the both triple negative samples and the natural thresholds set after clustering samples. Log2cpm for each gene; *ESR1*: 6, *PGR:* 6 and *ERBB2*: 10.5.

### Clustering of DCIS samples

Clustering and visualizations were done in R using all DCIS samples. The Limma-Voom ‘filterByExpr’ function was used to select genes expressed in at least 5% of the samples (*n*=19366). Raw counts were TMM-normalized and transformed into log_2_(CPM) values. To visualize the data and to reduce the variation driven by patient differences, we applied principal component analysis (PC analysis; PCA) using the ‘prcomp’ function with default settings. The number of PCs used in the subsequent clustering and UMAP steps was selected as the minimum number of PCs required to explain >30% of the total variance in the data (13 PCs). Hierarchical clustering was done using the ‘hclust’ function and the ward.D2 agglomeration method. The resulting tree was cut into five clusters, with triple negative samples forming 1 of the clusters. UMAP visualization was done using the ‘umap’ function from the umap package with default settings except increasing the number of epochs to 500, minimum distance to 0.2 and neighbours to 100 to reduce patient-specific effects.

### UMAP visualization of all samples

Visualization of all samples with UMAP was done in R. The Limma-Voom ‘filterByExpr’ function was used to select genes expressed in at least one of the tissue types (n=19661). Raw counts were TMM-normalized and transformed into log2(CPM) values. A PCA was constructed using the ‘prcomp’ function with default settings. UMAP visualization was done using the ‘umap’ function from the uwot package with default settings except setting the number of PCs to the minimum number of PCs required to explain >30% of the total variance in the data (16 PCs), increasing the number of epochs to 500, minimum distance to 0.2 and neighbours to 100 to reduce patient-specific effects

### Differential expression analysis

Differential expression analysis was done using Limma-Voom. First, expressed genes to include were selected by the ‘filterByExpr’ function using the design matrix as a guide, followed by calculation of normalization factors using the TMM method. To correct for the data structures with multiple samples coming from the same patient, we used a double ‘voom’ approach, including a ‘duplicateCorrection’ step with blocking based on patient. If no patient duplication was present in the contrast, we used a standard approach with a single application of ‘voom’. Fitting was done using ‘lmFit’ (with blocking and correction applied if applicable), followed by construction and calculation of contrasts using ‘contrast.fit’ function followed by ‘eBayes’. A gene was considered to be differentially expressed if the adjusted p-value was <0.05.

### Pseudo-time analysis

A differential expression analysis was done, as described above, between DCIS and IDC samples taken from the same patients, followed by a PCA using the most significant genes (p<0.00001, *n*=53). Remaining DCIS and IDC samples from patients without both types were projected onto this PCA embedding, together with samples from normal, benign and atypical epithelium. Since the different tissue types were positioned on the PCA in a biologically meaningful order, we fitted a principal curve to the data and projected the samples onto it to allow us to arrange them by their predicted pseudo-time order. We note that arranging them according to their position on a UMAP embedding resulted in largely the same order.

### Cell type enrichment analysis using xCell

Cell type enrichment scores were calculated using xCell (25) applied to FPKM-transformed counts. All studied samples (2222) were processed together. To study the change in epithelial cell enrichment across the timeline, we sorted the samples in timeline order and calculated a linear regression fit using the ‘lm’ function in R. Confidence intervals of the correlation coefficient r^2^ and the fitted line were estimated by bootstrapping using residual resampling with 10000 replicates.

### Gene set enrichment analysis

The R Bioconductor package RITAN (v.1.10.0) was used for gene set enrichment analysis using the MSigDB Hallmarks database All protein-coding genes were used as a background. Terms with FDR-adjusted p-value < 1e-5 are listed. To determine enrichment across the timeline, we used a sliding window of 100 samples, moving 50 samples at a time, compared to all remaining samples.

### Patient marker classifier

High and low expression was based on the majority segregation between Pure DCIS and Not Pure DCIS. Table S2 provides the expression values in log2 Counts per Million (CPM) for each marker.

A patient was placed in a group on the decision tree based on a minimum of 2 samples representing the “associated with IDC” expression levels, this being low *MNX1*, low *HOXC11*, low *ANKRD22*, low *ADCY5*, High *SCGB2A1*, low *CAMK2N1* and low *THRSP*. Two patients were removed from the decision trees as data was only available for 1 sample.

---

**Figure S1.**
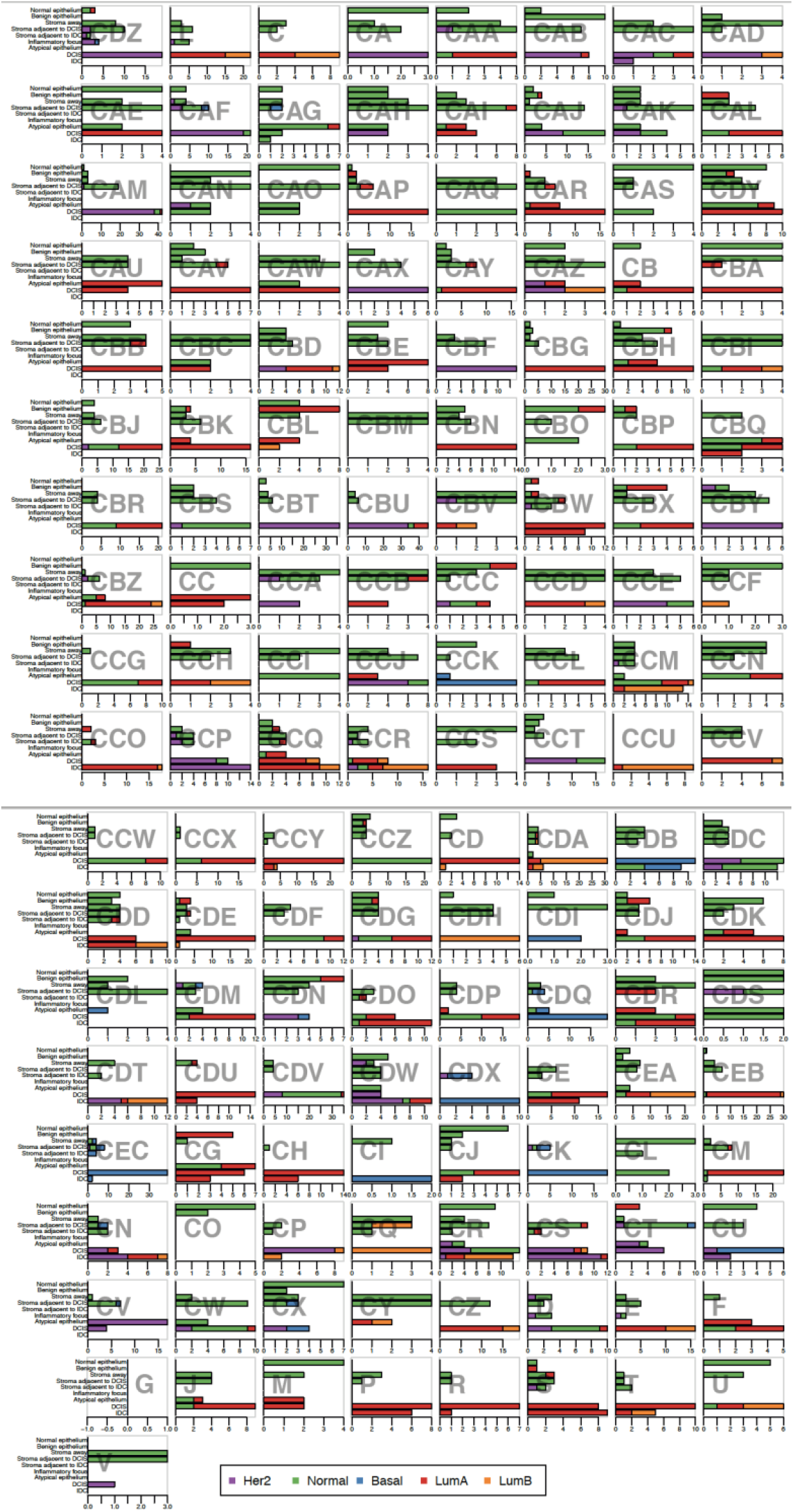
Patient subtype classification. Number of samples (after filtering) assigned with each AIMS subtype classification, from each patient. We found 52% of patients had mixed AIMS classifications for their DCIS samples, and 46% having mixed classifications for their IDC samples.

**Figure S2.**
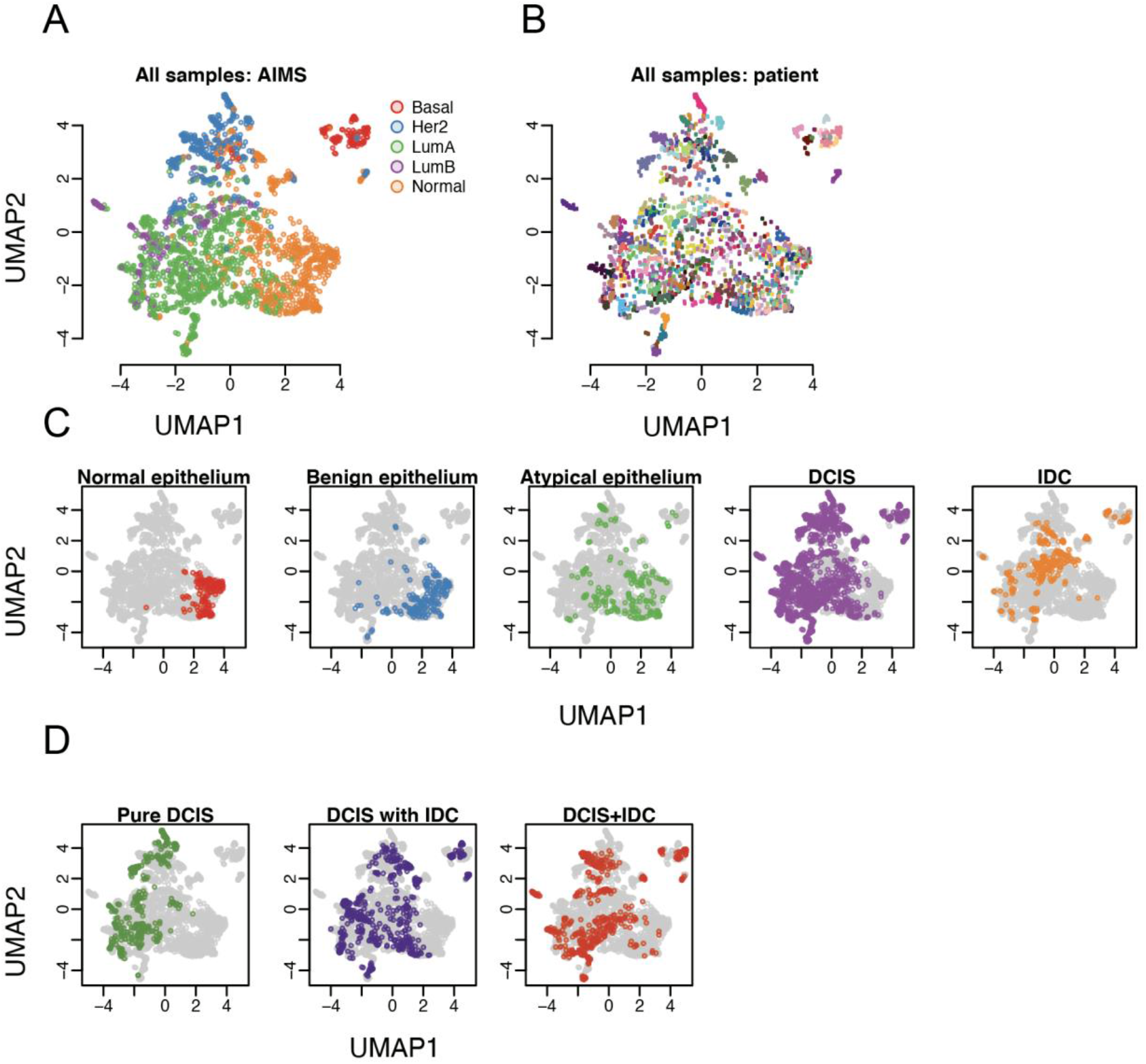
Sample clustering. Principal component analysis (PCA) followed by uniform manifold approximation and projection (UMAP) for all samples that passed quality filters. (A) All samples coloured by their AIMS subtype classification. (B) All samples coloured by which patient they came from. (C) Distribution of each tissue type – coloured - against all samples – grey. (D) Distribution of each DCIS group – coloured – against all samples – grey.

**Figure S3.**
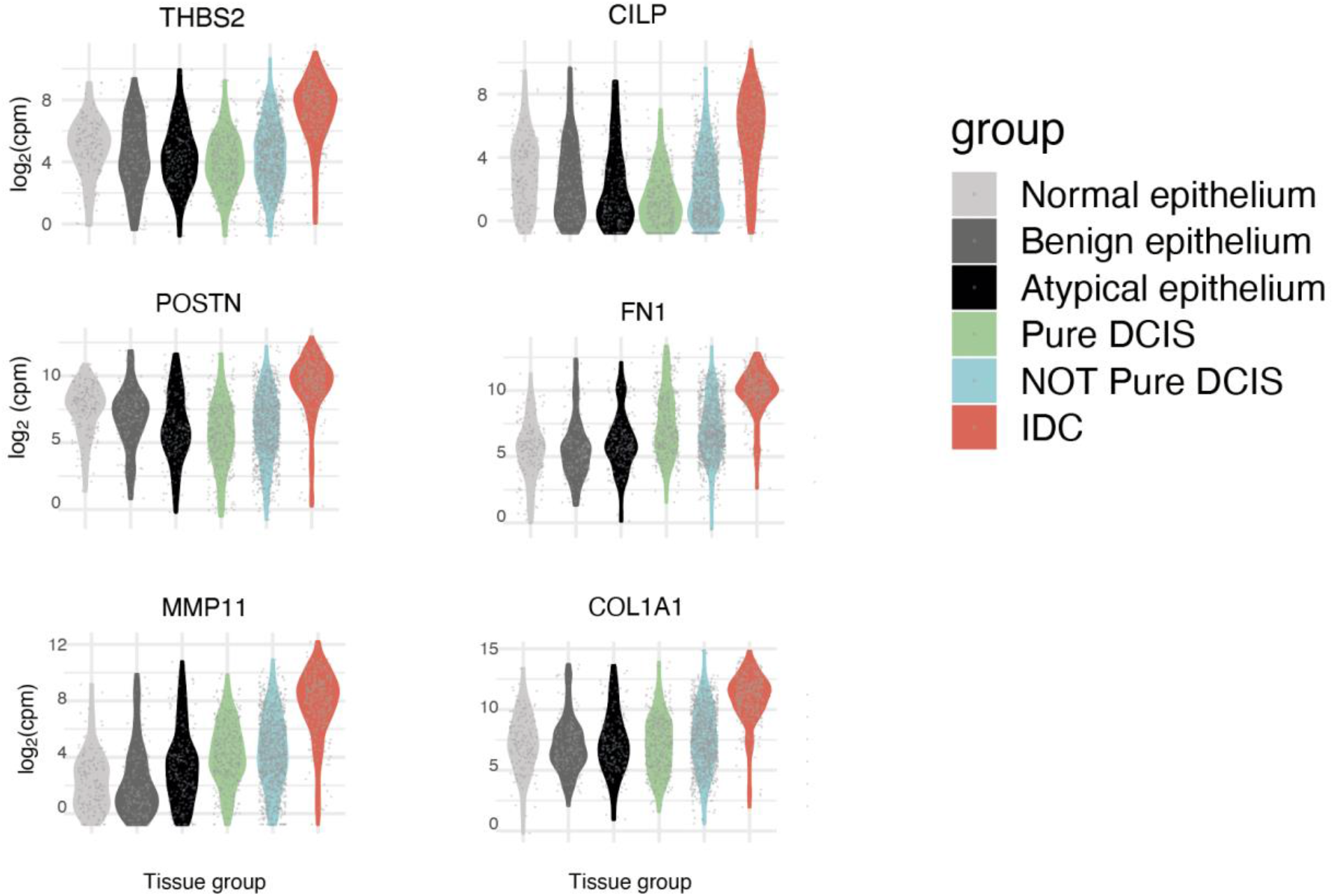
Differentially expressed genes between DCIS and co-occurring IDC. Expression distribution for example genes that showed a progressive shift among different tissue groups. Each sample is represented by a grey dot and a kernel density plot is over laid.

**Figure S4.**
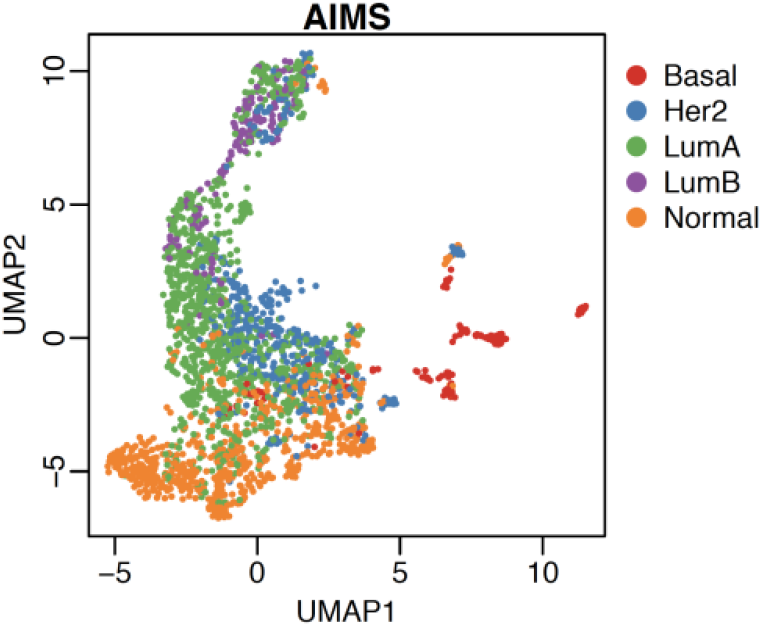
UMAP visualization using the same 53 genes that were used to construct the PCA plot in Fig. 3a. UMAP separates the samples more strongly by subtype compared with PCA and most triple-negative (basal) samples cluster separately.

**Figure S5.**
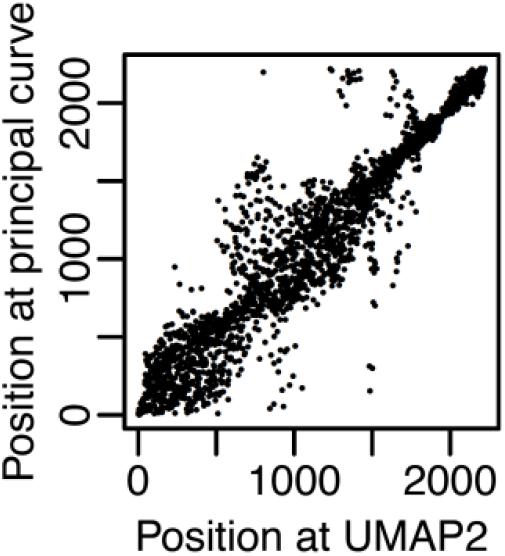
Ranking of samples across UMAP2 and across the principal curve, using the same 53 genes used to construct the PCA plot in Fig. 3a.

**Figure S6.**
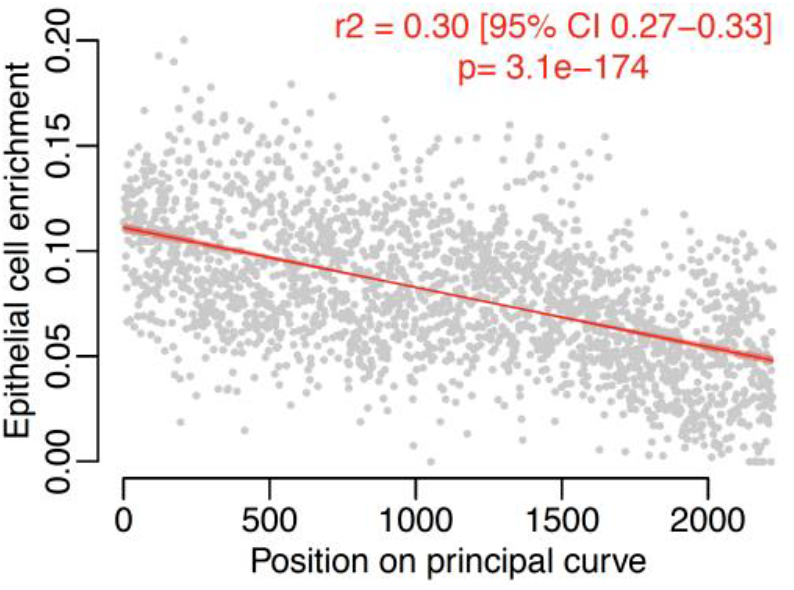
Epithelial cell enrichment calculated using xCell. All samples are sorted in timeline order. The line indicates the linear regression fit with a 95% confidence interval.

**Figure S7.**
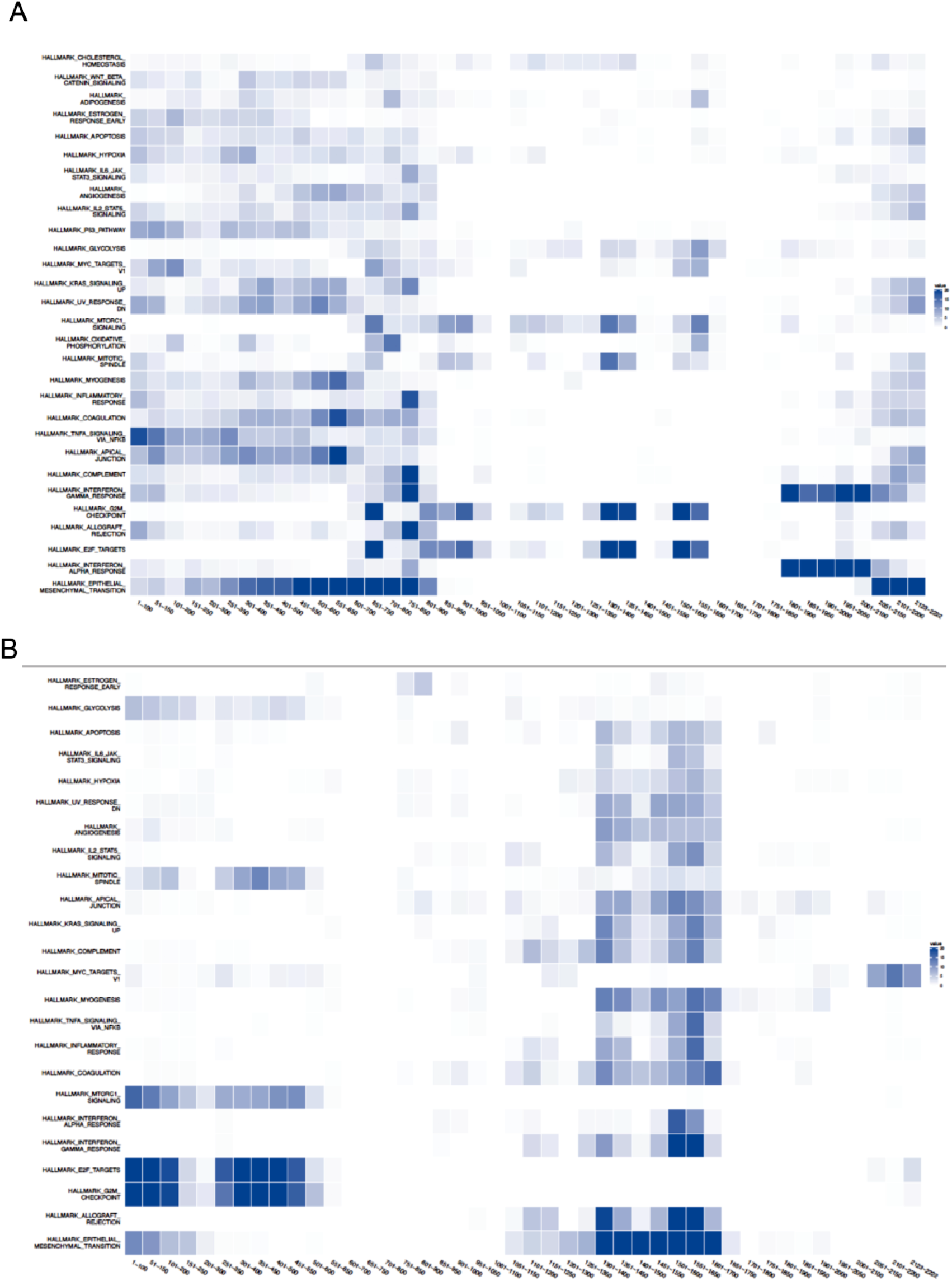
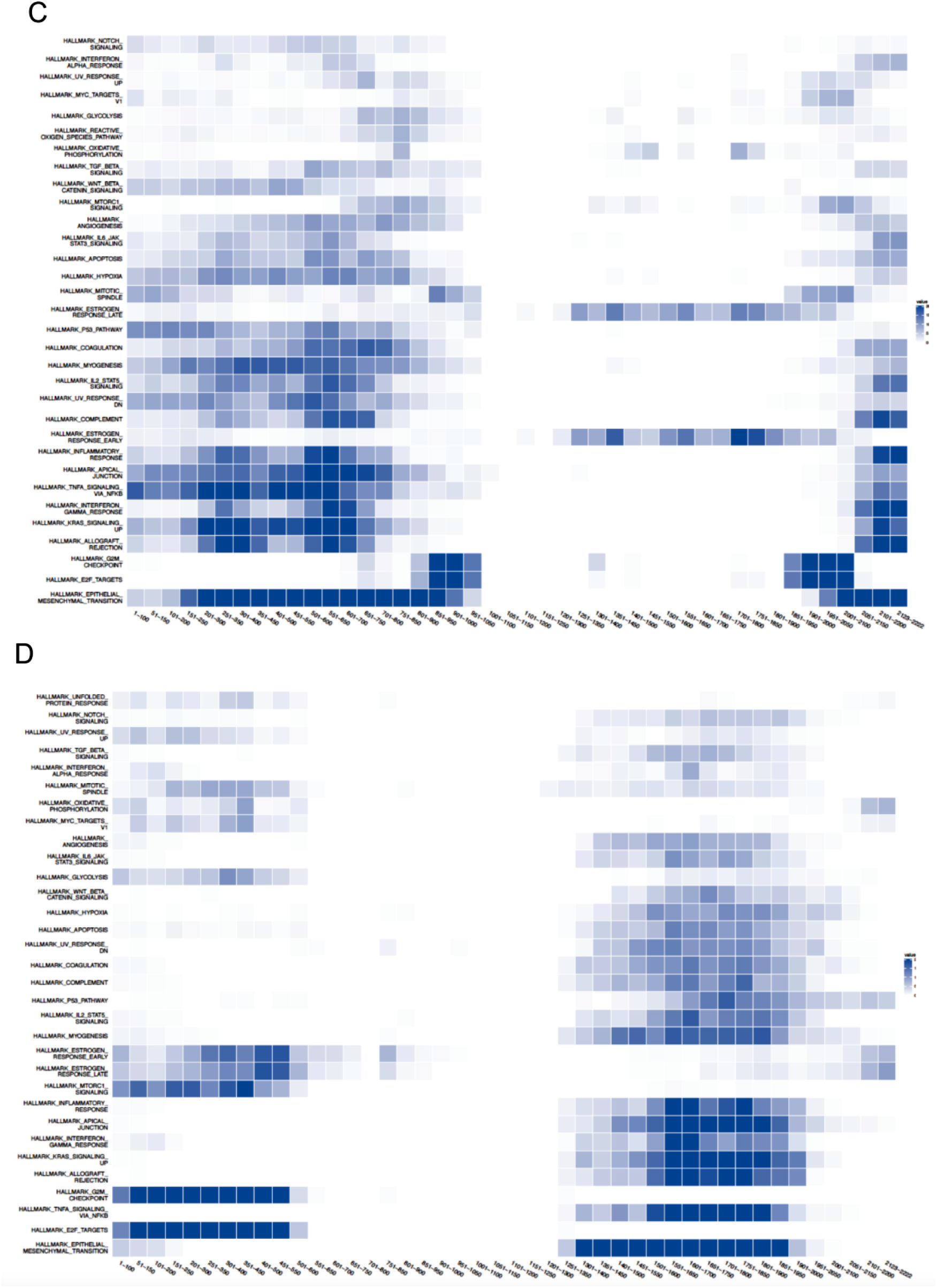
Related to Figure 4. Hallmark signatures up along the Timeline. Up (**A**) and down (**B**) for all ER positive samples. Up (**C**) and down (**D**) for all ER negative. Samples in order of Timeline. Sample average is calculated for each bin and compared to the average for all samples

**Figure S8.**
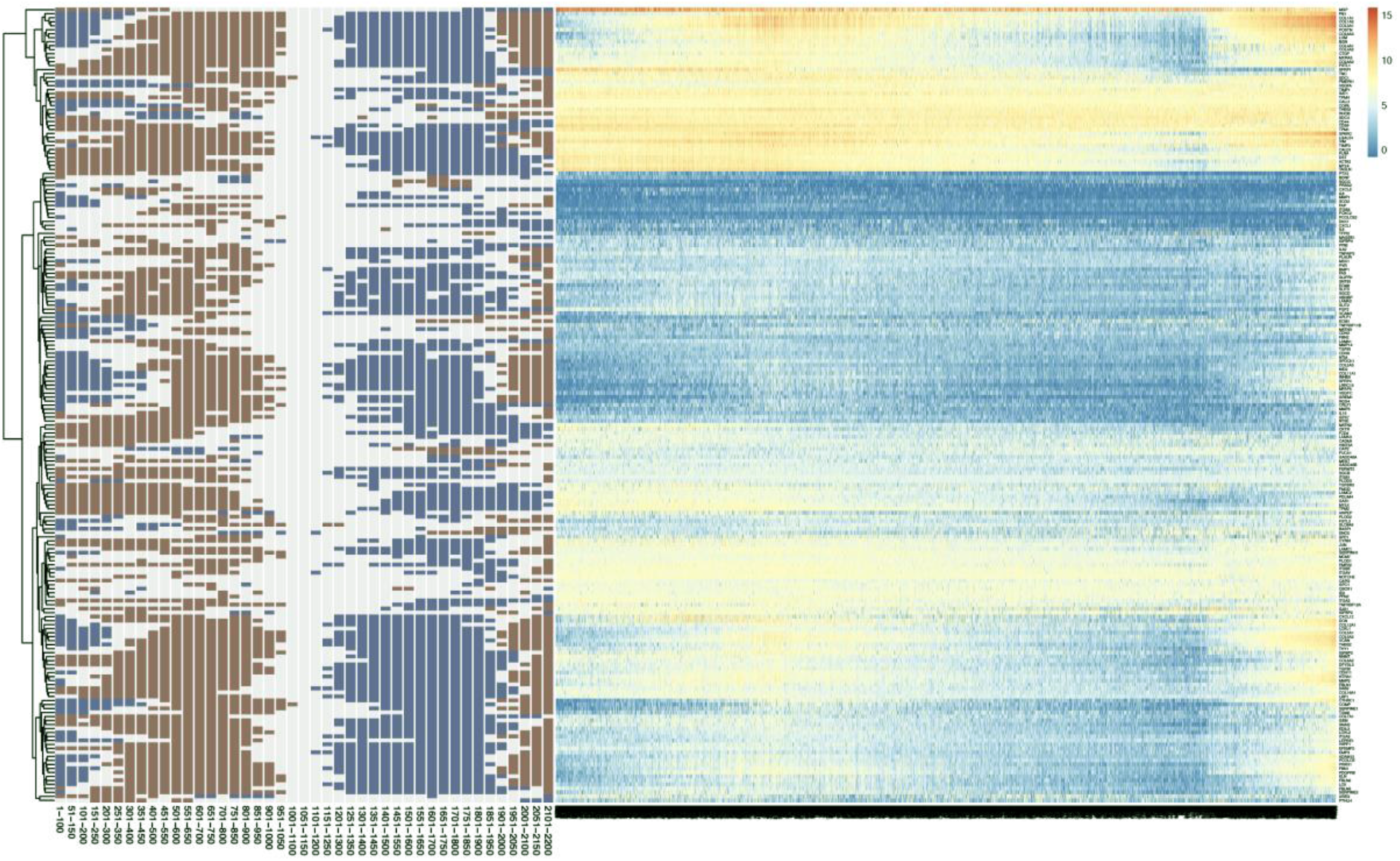
Epithelial to Mesenchymal transition occurs twice in the timeline. Heatmap showing relative gene expression for genes listed within the Epithelial to Mesenchymal transition Hallmark signature. Samples were ordered according to the projected principal curve. Bars to the left of the heat map reflect the differential expression analysis between the 100 samples in that block against all other samples. Genes that were significantly up- (red) or down-regulated (blue) are highlighted. The threshold for being red or blue was p.adj.<0.05.

**Figure S9.**
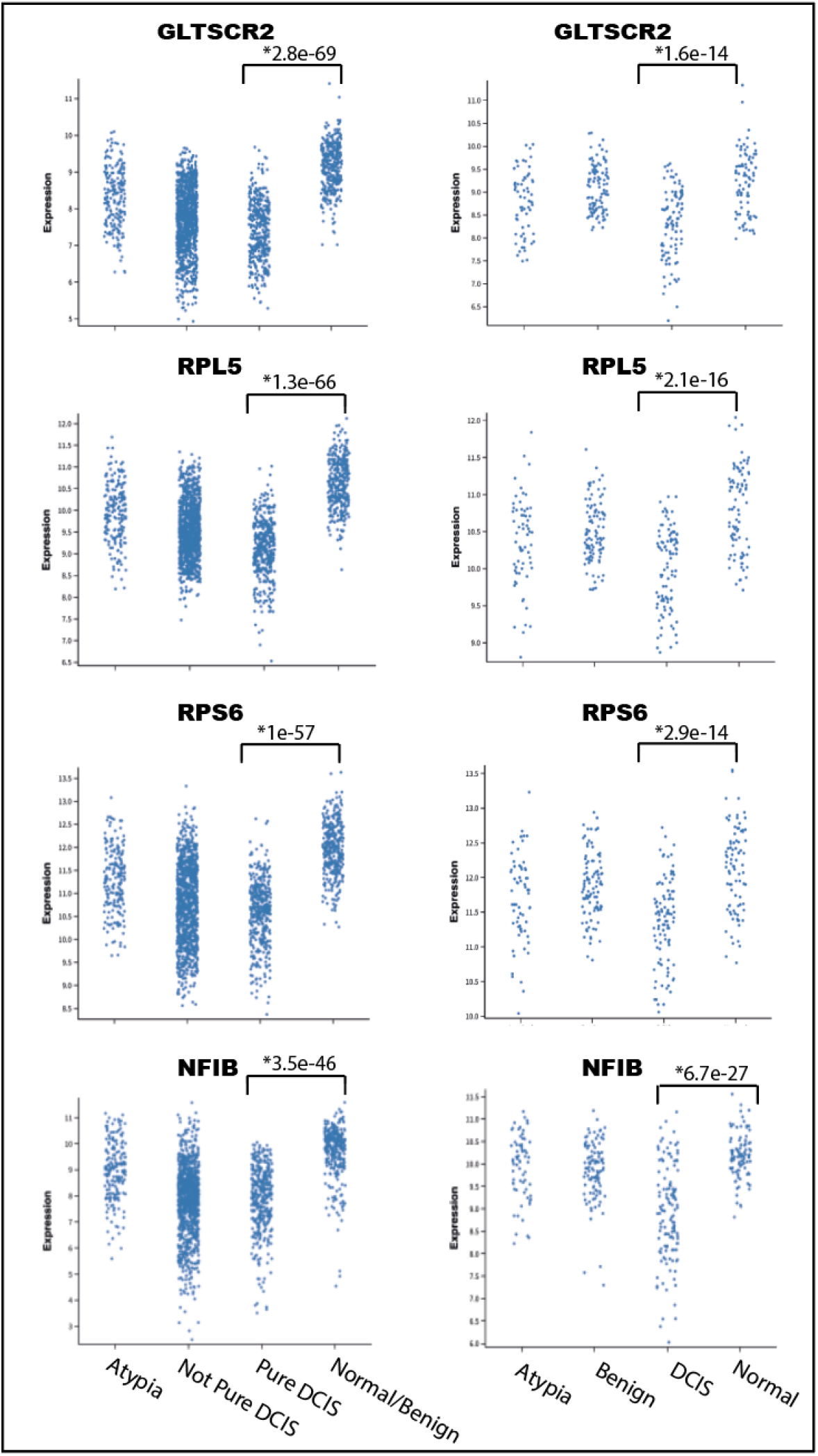
Differentially Expressed genes found when comparing normal/benign ductal tissue to DCIS samples. Expression distribution of samples in Log2 counts per million (CPM). Left panel show all samples with each tissue type, right panel show samples in the very early timeline. * indicates Adj. P value.

**Figure S10.**
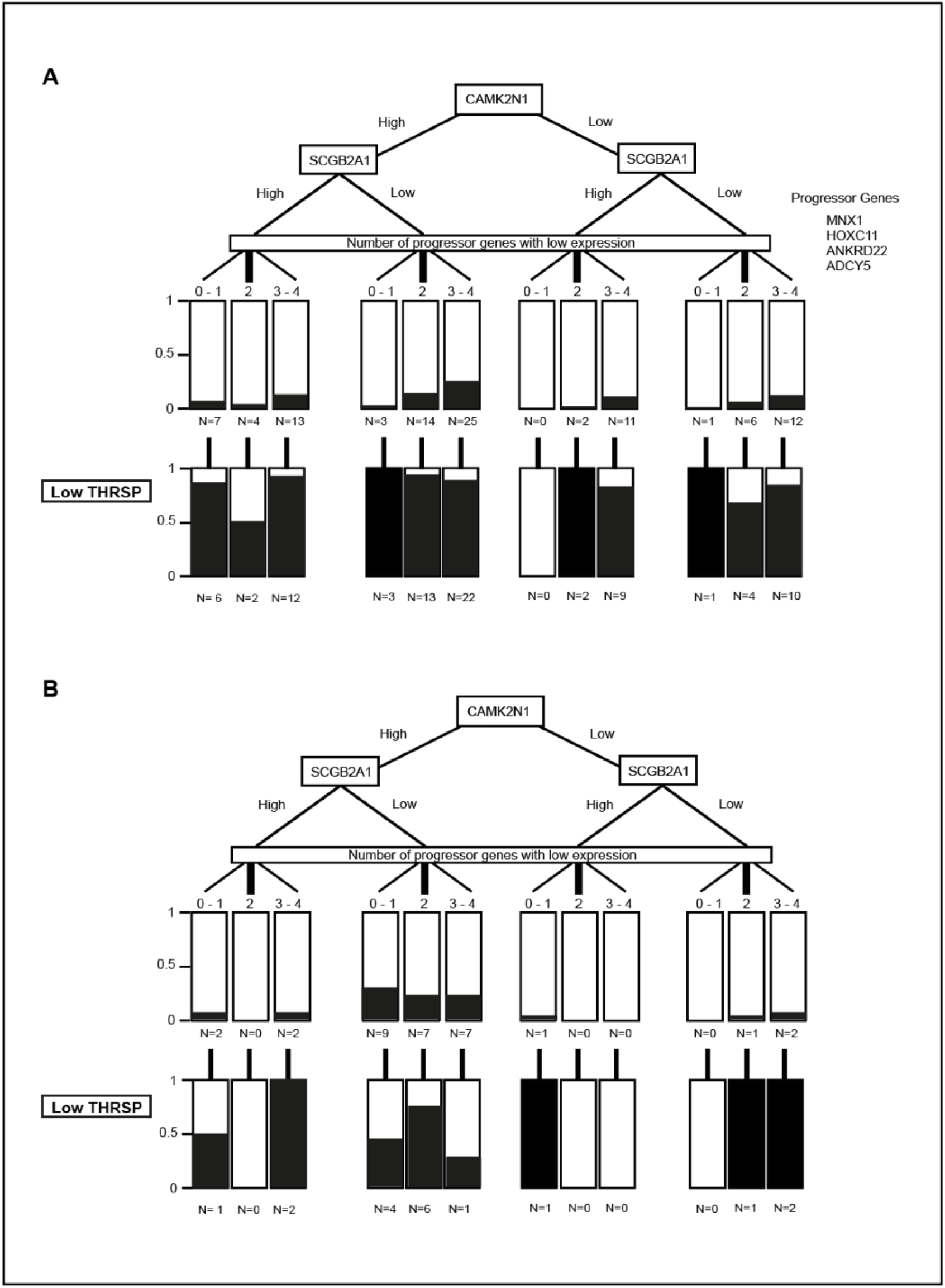
Decision tree for all samples. Separation of all patients **A**. diagnosed with IDC, N = 98 [2 patients excluded (*see supplementary methods*)], and **B**. that were never diagnosed with IDC (Pure DCIS), N = 31. Black bars represent the proportion of the total that fall in that node, e.g. N = 2 is 2% where the total is 98 or 6.4% where the total is 31. Boxes in the low *THRSP* layer represent the proportion of the group above.

**Table S1.**
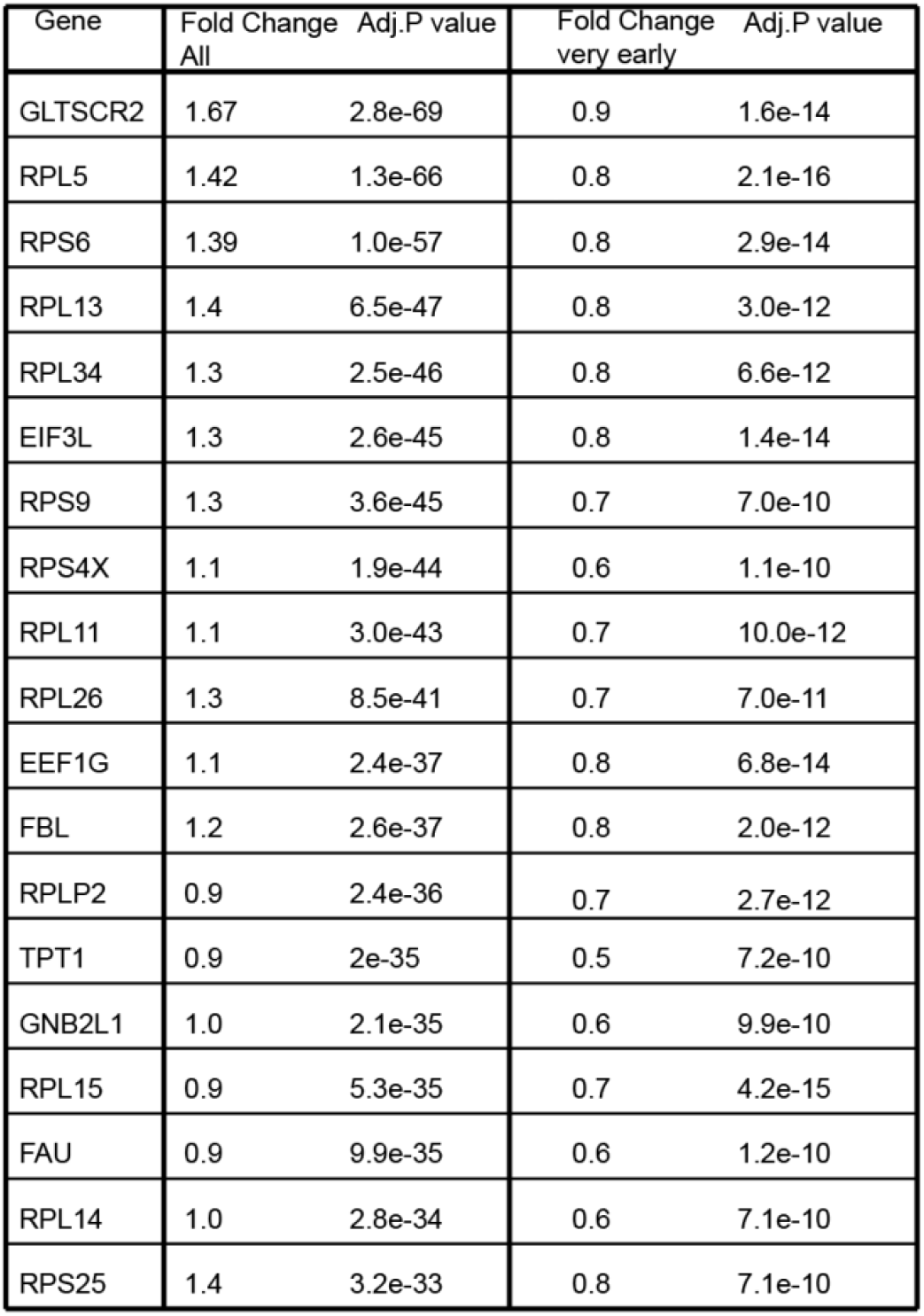
Ribosomal biogenesis genes significantly down regulated in DCIS compared to normal tissue. All – refers to analysis comparing all normal/benign tissues with Pure DCIS Very early – refers to analysis comparing normal tissues with DCIS tissues in the very early part of the timeline. Gene list represents the cluster of highly significant genes that were shared between All analysis and very early analysis

| Gene | Fold Change<br>All | Adj.P value | Fold Change<br>very early | Adj.P value |
| --- | --- | --- | --- | --- |
| GLTSCR2 | 1.67 | 2.8e-69 | 0.9 | 1.6e-14 |
| RPL5 | 1.42 | 1.3e-66 | 0.8 | 2.1e-16 |
| RPS6 | 1.39 | 1.0e-57 | 0.8 | 2.9e-14 |
| RPL13 | 1.4 | 6.5e-47 | 0.8 | 3.0e-12 |
| RPL34 | 1.3 | 2.5e-46 | 0.8 | 6.6e-12 |
| EIF3L | 1.3 | 2.6e-45 | 0.8 | 1.4e-14 |
| RPS9 | 1.3 | 3.6e-45 | 0.7 | 7.0e-10 |
| RPS4X | 1.1 | 1.9e-44 | 0.6 | 1.1e-10 |
| RPL11 | 1.1 | 3.0e-43 | 0.7 | 10.0e-12 |
| RPL26 | 1.3 | 8.5e-41 | 0.7 | 7.0e-11 |
| EEF1G | 1.1 | 2.4e-37 | 0.8 | 6.8e-14 |
| FBL | 1.2 | 2.6e-37 | 0.8 | 2.0e-12 |
| RPLP2 | 0.9 | 2.4e-36 | 0.7 | 2.7e-12 |
| TPT1 | 0.9 | 2e-35 | 0.5 | 7.2e-10 |
| GNB2L1 | 1.0 | 2.1e-35 | 0.6 | 9.9e-10 |
| RPL15 | 0.9 | 5.3e-35 | 0.7 | 4.2e-15 |
| FAU | 0.9 | 9.9e-35 | 0.6 | 1.2e-10 |
| RPL14 | 1.0 | 2.8e-34 | 0.6 | 7.1e-10 |
| RPS25 | 1.4 | 3.2e-33 | 0.8 | 7.1e-10 |

**Table S2.**
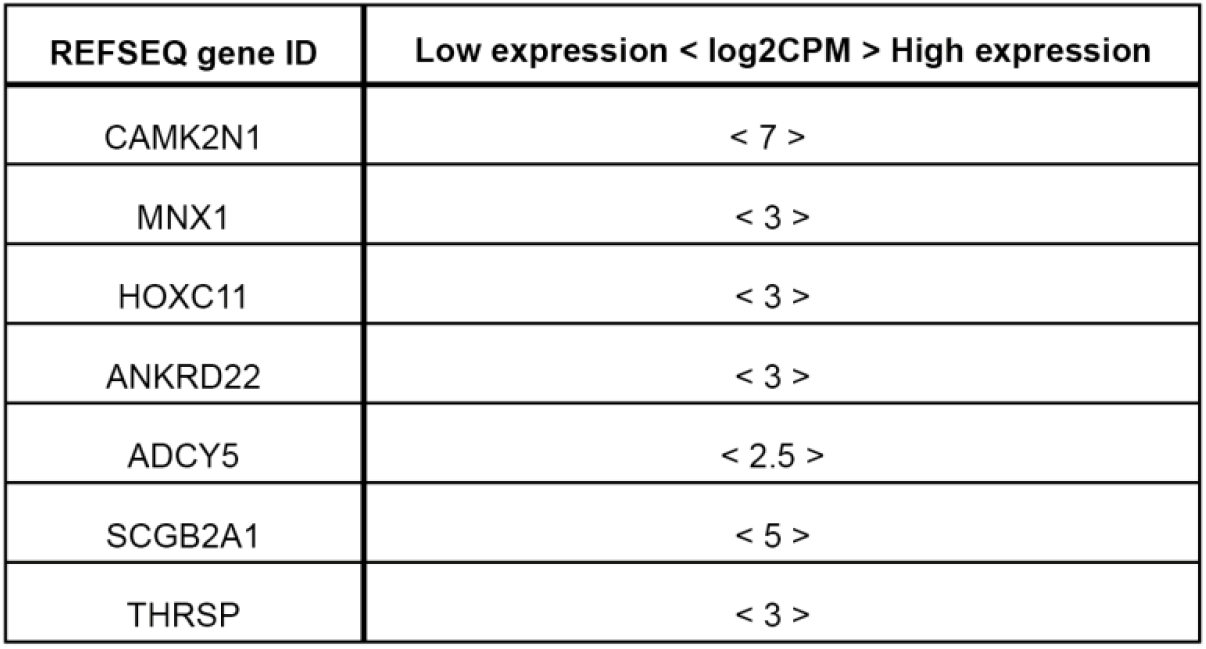
Gene expression thresholds. Distinction for high and low expression for each gene used in the classification (in log_2_ counts per million (CPM)).

**Table S3.** Differential genes in the High Hazard group. Genes distinguishing Pure DCIS from DCIS associated with IDC (Not Pure DCIS) in the Higher Hazard group of patients. Differential genes are from analysis first using only patients with *CAMK2N1* high / *SCGB2A1* low and reduced expression of 3-4 progressor genes, and then second using all patients with reduced expression of 3-4 progressor genes, regardless of *CAMK2N1* or *SCGB2A1* expression.

|  | CAMK2N1 + / SCGB2A1 - / 3-4 progressor genes down |  |  |  | All 3-4 progressor genes down |  |  |  |
| --- | --- | --- | --- | --- | --- | --- | --- | --- |
| REFSEQ gene ID | log2FC | Adj.PValue | Mean Expression |  | log2FC | Adj.PValue | Mean Expression |  |
|  |  |  | Pure DCIS | Not Pure DCIS |  |  | Pure DCIS | Not Pure DCIS |
| PHGR1 | 4.33 | 8.4e-13 | 7.88 | 4.00 | 4.07 | 3.05e-22 | 6.93 | 3.93 |
| THRSP | 4.04 | 1.5e-10 | 5.54 | 1.69 | 1.5 | 0.01 | 3.86 | 2.11 |
| SERPINA5 | 2.48 | 1e-8 | 7.00 | 4.74 | 2.8 | 7.24e-19 | 6.86 | 4.48 |
| LYPD6B | 1.36 | 0.01 | 6.4 | 5.11 | 2.45 | 3.28e-17 | 6.36 | 4.01 |
| GFRA1 | 1.88 | 9.3e-4 | 9.25 | 7.38 | 2.97 | 6.9e-17 | 9.12 | 5.99 |
| NPNT | 1.6 | 0.004 | 8.12 | 6.15 | 2.04 | 5.4e-10 | 7.85 | 5.38 |
| SLPI | 2.01 | 0.25 | 2.18 | 4.01 | 2.7 | 0.03 | 2.32 | 4.99 |
| SERPINE2 | 2.31 | 0.14 | 1.39 | 3.11 | 2.46 | 0.01 | 1.57 | 3.49 |
| FBLN2 | 2.22 | 0.1 | 1.11 | 2.92 | 2.43 | 0.006 | 1.23 | 3.23 |
| MSL3P1 | 1.51 | 0.27 | 0.05 | 1.4 | 2.17 | 0.004 | 0 | 1.95 |

## References and Notes

1. Mannu, G. S. et al. Invasive breast cancer and breast cancer mortality after ductal carcinoma in situ in women attending for breast screening in England, 1988-2014: population based observational cohort study. Bmj 369, m1570, doi:10.1136/bmj.m1570 (2020).

2. Collins, L. C. et al. Outcome of patients with ductal carcinoma in situ untreated after diagnostic biopsy: results from the Nurses’ Health Study. Cancer 103, 1778–1784, doi:10.1002/cncr.20979 (2005).

3. Welch, H. G. & Black, W. C. Using autopsy series to estimate the disease "reservoir" for ductal carcinoma in situ of the breast: how much more breast cancer can we find? Ann Intern Med 127, 1023–1028, doi:10.7326/0003-4819-127-11-199712010-00014 (1997).

4. Boecker, W. et al. Ductal epithelial proliferations of the breast: a biological continuum? Comparative genomic hybridization and high-molecular-weight cytokeratin expression patterns. J Pathol 195, 415–421, doi:10.1002/path.982 (2001).

5. Doebar, S. C. et al. Gene Expression Differences between Ductal Carcinoma in Situ with and without Progression to Invasive Breast Cancer. Am J Pathol 187, 1648–1655, doi:10.1016/j.ajpath.2017.03.012 (2017).

6. Kothari, C. et al. Identification of a gene signature for different stages of breast cancer development that could be used for early diagnosis and specific therapy. Oncotarget 9, 37407–37420, doi:10.18632/oncotarget.26448 (2018).

7. Dettogni, R. S. et al. Potential biomarkers of ductal carcinoma in situ progression. BMC Cancer 20, 119, doi:10.1186/s12885-020-6608-y (2020).

8. Gregory, K. J. et al. Gene expression signature of atypical breast hyperplasia and regulation by SFRP1. Breast Cancer Research 21, 76, doi:10.1186/s13058-019-1157-5 (2019).

9. Risom, T. et al. Transition to invasive breast cancer is associated with progressive changes in the structure and composition of tumor stroma. Cell 185, 299–310, doi:10.1016/j.cell.2021.12.023 (2022).

10. Paquet, E. R. & Hallett, M. T. Absolute assignment of breast cancer intrinsic molecular subtype. J Natl Cancer Inst 107, 357, doi:10.1093/jnci/dju357 (2015).

11. Bergholtz, H. et al. Contrasting DCIS and invasive breast cancer by subtype suggests basal- like DCIS as distinct lesions. NPJ Breast Cancer 6, 26, doi:10.1038/s41523-020-0167-x (2020).

12. Li, C. M.-C. et al. Aging-associated alterations in mammary epithelia and stroma revealed by single-cell RNA sequencing. Cell Reports 33, 108566, doi:https://doi.org/10.1016/j.celrep.2020.108566 (2020).

13. Dai, X. et al. FOXA1 is prognostic of triple negative breast cancers by transcriptionally suppressing SOD2 and IL6. Int J Biol Sci 15, 1030–1041, doi:10.7150/ijbs.31009 (2019).

14. G M Bernardo, G. B., C L Ginther, S T Sizemore, K L Lozada, J D Miedler, L A Anderson, A K Godwin, F W Abdul-Karim, D J Slamon, R A Keri. FOXA1 represses the molecular phenotype of basal breast cancer cells. Oncogene 32, 554–563, doi:10.1038/onc.2012.62 (2013).

15. Chivukula, M. et al. Prognostic significance of transcription factors FOXA1 and GATA-3 in ductal carcinoma *in situ* in terms of recurrence and estrogen receptor status. Journal of Cancer Metastasis and Treatment 1, 84–89, doi:10.4103/2394-4722.157600 (2015).

16. Picarsic, J., Brufsky, A., Ahrendt, G., Tseng, G. & Chivukula, M. Role of transcription factors [FOXA1,GATA-3] in predicting outcomes in recurrent ductal carcinoma-In-situ (DCIS) or invasive carcinoma (IC) in DCIS patients on core needle biopsies of breast. Cancer Research 69, 2115, doi:10.1158/0008-5472.SABCS-09-2115 (2009).

17. Segaert, P., Lopes, M. B., Casimiro, S., Vinga, S. & Rousseeuw, P. J. Robust identification of target genes and outliers in triple-negative breast cancer data. Stat Methods Med Res 28, 3042–3056, doi:10.1177/0962280218794722 (2019).

18. Cho, W. C., Ma, V. W., Cheuk, W., So, G. Y. & Chin, R. Y. Abstract 3148: FOXC1 expression is associated with a triple-negative basal-like phenotype in breast cancer. Cancer Research 80, 3148, doi:10.1158/1538-7445.AM2020-3148 (2020).

19. He, J. et al. Molecular features of triple negative breast cancer: microarray evidence and further integrated analysis. PloS one 10, e0129842–e0129842, doi:10.1371/journal.pone.0129842 (2015).

20. Szklarczyk, D. et al. STRING v11: protein-protein association networks with increased coverage, supporting functional discovery in genome-wide experimental datasets. Nucleic Acids Res 47, D607–d613, doi:10.1093/nar/gky1131 (2019).

21. Caubet, C. et al. Degradation of corneodesmosome proteins by two serine proteases of the kallikrein family, SCTE/KLK5/hK5 and SCCE/KLK7/hK7. J Invest Dermatol 122, 1235–1244, doi:10.1111/j.0022-202X.2004.22512.x (2004).

22. Pal, B. et al. Construction of developmental lineage relationships in the mouse mammary gland by single-cell RNA profiling. Nature communications 8, 1627–1627, doi:10.1038/s41467-017-01560-x (2017).

23. Huper, G. & Marks, J. R. Isogenic normal basal and luminal mammary epithelial isolated by a novel method show a differential response to ionizing radiation. Cancer Research 67, 2990, doi:10.1158/0008-5472.CAN-06-4065 (2007).

24. Aran, D., Hu, Z. & Butte, A. J. xCell: digitally portraying the tissue cellular heterogeneity landscape. Genome Biol 18, 220, doi:10.1186/s13059-017-1349-1 (2017).

25. Karamanou, K. et al. Lumican effectively regulates the estrogen receptors-associated functional properties of breast cancer cells, expression of matrix effectors and epithelial-to-mesenchymal transition. Sci Rep 7, 45138, doi:10.1038/srep45138 (2017).

26. Stevenson, A. J. et al. Multiscale imaging of basal cell dynamics in the functionally mature mammary gland. Proc Natl Acad Sci U S A 117, 26822–26832, doi:10.1073/pnas.2016905117 (2020).

27. Du, Y. et al. The cancer-associated fibroblasts related gene CALD1 is a prognostic biomarker and correlated with immune infiltration in bladder cancer. Cancer Cell Int 21, 283, doi:10.1186/s12935-021-01896-x (2021).

28. Zheng, P. P. et al. Differential expression of splicing variants of the human caldesmon gene (CALD1) in glioma neovascularization versus normal brain microvasculature. Am J Pathol 164, 2217–2228, doi:10.1016/S0002-9440(10)63778-9 (2004).

29. Russell, T. D. et al. Myoepithelial cell differentiation markers in ductal carcinoma in situ progression. Am J Pathol 185, 3076–3089, doi:10.1016/j.ajpath.2015.07.004 (2015).

30. Casasent, A. K. et al. Multiclonal Invasion in Breast Tumors Identified by Topographic Single Cell Sequencing. Cell 172, 205–217.e212, doi:10.1016/j.cell.2017.12.007 (2018).

31. Rakovitch, E. et al. Significance of multifocality in ductal carcinoma in situ: outcomes of women treated with breast-conserving therapy. J Clin Oncol 25, 5591–5596, doi:10.1200/jco.2007.11.4686 (2007).

32. Whitfield, M. L., George, L. K., Grant, G. D. & Perou, C. M. Common markers of proliferation. Nature Reviews Cancer 6, 99–106, doi:10.1038/nrc1802 (2006).

33. Kim, J.-Y. et al. Involvement of GLTSCR2 in the DNA damage response. The American journal of pathology 179, 1257–1264, doi:10.1016/j.ajpath.2011.05.041 (2011).

34. Kim, Y. J. et al. Suppression of putative tumour suppressor gene GLTSCR2 expression in human glioblastomas. J Pathol 216, 218–224, doi:10.1002/path.2401 (2008).

35. Lee, S. et al. Nucleolar protein GLTSCR2 stabilizes p53 in response to ribosomal stresses. Cell Death Differ 19, 1613–1622, doi:10.1038/cdd.2012.40 (2012).

36. Fumagalli, S., Ivanenkov, V. V., Teng, T. & Thomas, G. Suprainduction of p53 by disruption of 40S and 60S ribosome biogenesis leads to the activation of a novel G2/M checkpoint. Genes & development 26, 1028–1040, doi:10.1101/gad.189951.112 (2012).

37. Amsterdam, A. et al. Many ribosomal protein genes are cancer genes in zebrafish. PLoS Biol 2, E139, doi:10.1371/journal.pbio.0020139 (2004).

38. Morgado-Palacin, L. et al. Partial Loss of Rpl11 in Adult Mice Recapitulates Diamond-Blackfan Anemia and Promotes Lymphomagenesis. Cell Rep 13, 712–722, doi:10.1016/j.celrep.2015.09.038 (2015).

39. Isidoro Cobo, S. P., Júlia Melià-Alomà, Ariadna Torres, Jaime Martínez-Villarreal, Fernando García, Irene Millán, Natalia del Pozo, Joo-Cheol Park, Ray J. MacDonald, Javier Muñoz, ProfileFrancisco X. Real. NFIC regulates ribosomal biology and ER stress in pancreatic acinar cells and suppresses PDAC initiation (2021).

40. Zilli, F. et al. The NFIB-ERO1A axis promotes breast cancer metastatic colonization of disseminated tumour cells. EMBO Mol Med 13, e13162, doi:10.15252/emmm.202013162 (2021).

41. Denny, S. K. et al. Nfib Promotes Metastasis through a Widespread Increase in Chromatin Accessibility. Cell 166, 328–342, doi:10.1016/j.cell.2016.05.052 (2016).

42. Grabowska, M. M. et al. Nfib Regulates Transcriptional Networks That Control the Development of Prostatic Hyperplasia. Endocrinology 157, 1094–1109, doi:10.1210/en.2015-1312 (2016).

43. Risom, T. et al. Transition to invasive breast cancer is associated with progressive changes in the structure and composition of tumor stroma. bioRxiv, 2021.2001.2005.425362, doi:10.1101/2021.01.05.425362 (2021).

44. Chi, M. et al. Phosphorylation of calcium/calmodulin-stimulated protein kinase II at T286 enhances invasion and migration of human breast cancer cells. Scientific Reports 6, 33132, doi:10.1038/srep33132 (2016).

45. Heinze, K. et al. CAMK2N1/RUNX3 methylation is an independent prognostic biomarker for progression-free and overall survival of platinum-sensitive epithelial ovarian cancer patients. Clinical Epigenetics 13, 15, doi:10.1186/s13148-021-01006-8 (2021).

46. Wang, T. et al. The tumor suppressive role of CAMK2N1 in castration-resistant prostate cancer. Oncotarget 5, 3611–3621, doi:10.18632/oncotarget.1968 (2014).

47. Bao, D. et al. Overexpression of CAMK 2 N 1 indicates good prognosis for glioma and regulates androgen receptor-associated cell proliferation and apoptosis. Int J Clin Exp Med 12, 540–548 (2019).

48. Butner, J. D. et al. A multiscale agent-based model of ductal carcinoma in situ. IEEE Trans Biomed Eng 67, 1450–1461, doi:10.1109/tbme.2019.2938485 (2020).

49. Cui, Y. et al. HOXC11 functions as a novel oncogene in human colon adenocarcinoma and kidney renal clear cell carcinoma. Life Sci 243, 117230, doi:10.1016/j.lfs.2019.117230 (2020).

50. Dai, B. W. et al. HOXC10 promotes migration and invasion via the WNT-EMT signaling pathway in oral squamous cell carcinoma. J Cancer 10, 4540–4551, doi:10.7150/jca.30645 (2019).

51. Kim, J. et al. HOXC10 overexpression promotes cell proliferation and migration in gastric cancer. Oncol Rep 42, 202–212, doi:10.3892/or.2019.7164 (2019).

52. Xiao, L., Hong, L. & Zheng, W. Motor neuron and pancreas homeobox 1 (MNX1) Is involved in promoting squamous cervical cancer proliferation via regulating cyclin E. Medical science monitor : international medical journal of experimental and clinical research 25, 6304–6312, doi:10.12659/MSM.914233 (2019).

53. Wu, Y., Liu, H., Gong, Y., Zhang, B. & Chen, W. ANKRD22 enhances breast cancer cell malignancy by activating the Wnt/β-catenin pathway via modulating NuSAP1 expression. Bosnian Journal of Basic Medical Sciences 21, 294–304, doi:10.17305/bjbms.2020.4701 (2021).

54. Yin, J. et al. ANKRD22 promotes progression of non-small cell lung cancer through transcriptional up-regulation of E2F1. Scientific reports 7, 4430–4430, doi:10.1038/s41598-017-04818-y (2017).

55. Qiu, Y. et al. ANKRD22 is involved in the progression of prostate cancer. Oncol Lett 18, 4106–4113, doi:10.3892/ol.2019.10738 (2019).

56. Sheng, H., Li, X. & Xu, Y. Knockdown of FOXP1 promotes the development of lung adenocarcinoma. Cancer biology & therapy 20, 537–545, doi:10.1080/15384047.2018.1537999 (2019).

57. Abba, M. C. et al. A Molecular Portrait of High-Grade Ductal Carcinoma In Situ. Cancer Res 75, 3980–3990, doi:10.1158/0008-5472.CAN-15-0506 (2015).

58. Tang, Q. & Hann, S. S. HOTAIR: An Oncogenic Long Non-Coding RNA in Human Cancer. Cell Physiol Biochem 47, 893–913, doi:10.1159/000490131 (2018).

59. Barrio, A. V. & Van Zee, K. J. Controversies in the Treatment of Ductal Carcinoma in Situ. Annual review of medicine 68, 197–211, doi:10.1146/annurev-med-050715-104920 (2017).

60. Gorringe, K. L. & Fox, S. B. Ductal carcinoma in situ biology, biomarkers, and diagnosis. Frontiers in oncology 7, 248–248, doi:10.3389/fonc.2017.00248 (2017).

61. Carter, D. et al. Purification and characterization of the mammaglobin/lipophilin B complex, a promising diagnostic marker for breast cancer. Biochemistry 41, 6714–6722, doi:10.1021/bi0159884 (2002).

62. Wellberg, E. A. et al. Modulation of tumor fatty acids, through overexpression or loss of thyroid hormone responsive protein spot 14 is associated with altered growth and metastasis. Breast Cancer Res 16, 481–481, doi:10.1186/s13058-014-0481-z (2014).

63. Namba, R. et al. Molecular characterization of the transition to malignancy in a genetically engineered mouse-based model of ductal carcinoma in situ. Mol Cancer Res 2, 453–463 (2004).

64. Asanuma, K. et al. Protein C inhibitor inhibits breast cancer cell growth, metastasis and angiogenesis independently of its protease inhibitory activity. Int J Cancer 121, 955–965, doi:10.1002/ijc.22773 (2007).

65. Bijsmans, I. T. et al. Loss of SerpinA5 protein expression is associated with advanced-stage serous ovarian tumors. Mod Pathol 24, 463–470, doi:10.1038/modpathol.2010.214 (2011).

66. Wagenblast, E. et al. A model of breast cancer heterogeneity reveals vascular mimicry as a driver of metastasis. Nature 520, 358–362, doi:10.1038/nature14403 (2015).

67. Erbas, B., Provenzano, E., Armes, J. & Gertig, D. The natural history of ductal carcinoma in situ of the breast: a review. Breast Cancer Res Treat 97, 135–144, doi:10.1007/s10549-005-9101-z (2006).

68. Sanders, M. E., Schuyler, P. A., Simpson, J. F., Page, D. L. & Dupont, W. D. Continued observation of the natural history of low-grade ductal carcinoma in situ reaffirms proclivity for local recurrence even after more than 30 years of follow-up. Mod Pathol 28, 662–669, doi:10.1038/modpathol.2014.141 (2015).

69. Saelens, W., Cannoodt, R., Todorov, H. & Saeys, Y. A comparison of single-cell trajectory inference methods. Nat Biotechnol 37, 547–554, doi:10.1038/s41587-019-0071-9 (2019).

70. Hanzelmann, S., Castelo, R. & Guinney, J. GSVA: gene set variation analysis for microarray and RNA-seq data. BMC Bioinformatics 14, 7, doi:10.1186/1471-2105-14-7 (2013).

71. Gonzalez-Solares, E. et al. The Imaging and Molecular Annotation of Xenografts and Tumours (IMAXT) High Throughput Data and Analysis Infrastructure. BioRxiv, doi:doi.org/10.1101/2021.06.22.448403 (2021).

72. Dobin, A. et al. STAR: ultrafast universal RNA-seq aligner. Bioinformatics 29, 15–21, doi:10.1093/bioinformatics/bts635 (2013).

73. Law, C. W., Chen, Y., Shi, W. & Smyth, G. K. voom: Precision weights unlock linear model analysis tools for RNA-seq read counts. Genome Biol 15, R29, doi:10.1186/gb-2014-15-2-r29 (2014).

